# Allosteric SHP2 Inhibition Increases Apoptotic Dependency on BCL2 and Synergizes with Venetoclax in *FLT3-* and *KIT-* Mutant AML

**DOI:** 10.1101/2022.12.01.518665

**Authors:** Bogdan Popescu, Carlos Stahlhut, Theodore C. Tarver, Sydney Wishner, Bianca J. Lee, Cheryl A. C. Peretz, Cuyler Luck, Paul Phojanakong, Juan Antonio Camara Serrano, Henry Hongo, Jose M. Rivera, Simayijiang Xirenayi, John A. Chukinas, Veronica Steri, Sarah K. Tasian, Elliot Stieglitz, Catherine C. Smith

## Abstract

Mutations in receptor tyrosine kinases (RTKs) FLT3 and KIT are frequent and associated with poor outcomes in acute myeloid leukemia (AML). Although FLT3 inhibitors (FLT3i) are clinically effective, remissions are short-lived due to secondary resistance characterized by acquired mutations constitutively activating the RAS/MAPK pathway. Hereby, we report pre-clinical efficacy of co-targeting SHP2, a critical node in MAPK signaling, and BCL2 in RTK-driven AML. The allosteric SHP2 inhibitor RMC-4550 suppressed proliferation of AML cell lines with FLT3 and KIT mutations, including cell lines with acquired resistance to FLT3i. We demonstrate that SHP2 inhibition unveils an Achilles’ heel of AML, increasing apoptotic dependency on BCL2 via MAPK-dependent mechanisms, including upregulation of BMF and downregulation of MCL1. Consequently, RMC-4550 and venetoclax are synergistically lethal in *FLT3*- or *KIT*-mutant AML cell lines, and in clinically relevant xenograft models. Our results provide new mechanistic rationale and preclinical evidence for co-targeting SHP2 and BCL2 in RTK-driven AML.

**Significance:** There is an unmet need for effective therapies targeting the MAPK pathway to overcome resistance in RTK-driven AML. We report that pharmacologic co-inhibition of SHP2 and BCL2 has synergistic anti-leukemia activity in preclinical models of AML with FLT3 and KIT mutations and holds potential clinical utility.

## Introduction

Mutations in the receptor tyrosine kinases (RTKs) encoded by *FLT3* and *KIT* are consistently among the most commonly identified molecular lesions in AML. These mutations occur in 30-40% of AML patients (1–3) and are associated with an increased risk of relapse. Patients with *FLT3* internal tandem duplication (ITD) mutations have decreased overall survival, primarily due to high relapse rates (4). Mutations in *KIT*, though more infrequent, are enriched in patients with AML driven by *CBFB-MYH11* or *RUNX1-RUNX1T1* gene fusions, and double the risk of relapse in a subtype of AML otherwise associated with good prognosis (5–6). Addition of the KIT inhibitor (KITi) dasatinib to chemotherapy in patients with newly-diagnosed core binding factor (CBF) AML is feasible (7–8), but improvements in relapse-free and overall survival have not been demonstrated in randomized trials. In contrast, FLT3 inhibitors (FLT3i) have proven benefit in AML patients with *FLT3* mutations in both upfront and relapsed/refractory (R/R) settings. Addition of the FLT3i midostaurin to standard induction chemotherapy demonstrated a survival benefit over chemotherapy alone in newly-diagnosed *FLT3*-mutant AML (9) and is currently the preferred first-line treatment in these patients. Second-generation selective FLT3 inhibitors quizartinib (AC220) and gilteritinib (ASP2215) showed superior overall survival compared to salvage chemotherapy in relapsed/refractory (R/R) *FLT3*-mutant AML in multinational phase 3 trials (10–11), establishing a new standard of care. However, remissions induced by these targeted therapies are short-lived and relapse due to secondary resistance limits clinical benefit (12–13). Given poor outcomes and the lack of consensus-validated standard of care for patients who relapse on FLT3i, there is a critical unmet need for novel treatment strategies to overcome clinical resistance.

Our group and others recently identified the emergence of polyclonal RAS/MAPK pathway mutations, most commonly in the form of activating *NRAS* mutations, in patients relapsing on FLT3i (13–15). Functional and biochemical analysis of AML cell lines implicate re-activation of MAPK signaling as a primary mediator of acquired resistance to FLT3i (13). *In vitro* modeling studies highlight the importance of protective bone marrow microenvironment cytokines in the progression of the MAPK-driven resistance. Specifically, soluble factors such as FLT3 ligand (FLT3-L) and fibroblast growth factor 2 (FGF2) reactivate ERK phosphorylation in residual AML cells and lay the groundwork for overt resistance, characterized by acquisition of intrinsic resistance mutations (16–17).

*RAS* pathway mutations have also been associated with intrinsic and adaptive resistance to other targeted AML therapies, including the IDH2 inhibitor enasidenib (18) and the BCL2 inhibitor venetoclax (19). While no studies have investigated the clinical resistance to KITi in AML, *RAS* mutations have been described in resistance to KITi in patients with KIT-mutant gastrointestinal stromal tumor (GIST, ref. 20). The body of evidence associating activating *RAS* mutations with clinical resistance to the full spectrum of approved targeted therapeutics in AML identifies a common and critical barrier to improved outcomes in AML.

Tyrosine-protein phosphatase non-receptor 11 (SHP2, encoded by *PTPN11*) is a protein tyrosine phosphatase (PTP) that functions as a signal relay molecule for RTK-mediated activation of the RAS/MAPK signaling pathway. Acting downstream of RTKs, SHP2 serves as a scaffold for the assembly of multi-protein complexes at the cell membrane that promote GTP loading of RAS by the guanine nucleotide exchange factor (GEF) SOS1 (21). SHP2 also promotes RAS activation by de-phosphorylating and inhibiting key RAS negative regulators such as RAS GTPase activating protein (RasGAP) (22) and Sprouty proteins (23). SHP2 has also been linked to RTK-dependent activation of phosphatidylinositol 3-kinase (PI3K)/AKT (24) and JAK/STAT signaling (25), and to upregulation of anti-apoptotic gene expression in FLT3-ITD+ cells (26–27). New allosteric SHP2 inhibitors (SHP2i) stabilize the closed, autoinhibited conformation, hampering both the catalytic and scaffolding functions of SHP2 (21, 28). Such compounds have demonstrated pre-clinical anti-tumor activity in multiple RTK-driven cancers, including *FLT3*-mutant AML *in vitro* and murine models (28–30). Moreover, allosteric SHP2i have shown activity in cell lines bearing RAS mutant proteins with substitutions at glycine 12 that remain dependent on GEF-mediated GTP loading for survival signaling (21, 31). SHP2i also prevents adaptive resistance to MEK inhibitors in both wild type and mutant KRAS cancer models (32).

Given the pivotal role of SHP2 in RTK signaling, we hypothesize that SHP2 inhibition will have activity in multiple subtypes of RTK-driven AML and will suppress key mechanisms of tyrosine kinase inhibitor (TKI) resistance dependent on activated MAPK signaling, particularly those driven by GTP-cycling RAS mutations and cytokine signaling. Although clinical-grade compounds such as TNO-155, RMC-4630, or JAB-3312 are under investigation in phase 1/2 clinical trials in a myriad of solid malignancies (33), SHP2 inhibition has not yet been clinically explored in AML, and effective targets for potential clinical combinations remain unknown.

Here, we show that the allosteric SHP2 inhibitor preclinical tool compound RMC-4550 (21) potently and selectively suppresses cell proliferation and induces apoptosis in *FLT3-* and *KIT-* mutant AML cell lines. Using high throughput transcriptomics, dynamic BH3 profiling and biochemical analyses, we demonstrate that SHP2 inhibition exposes an Achilles’ heel of AML cells by increasing apoptotic dependency on BCL2 via multiple MAPK-dependent mechanisms. Exploiting this vulnerability, we found that combining RMC-4550 and the FDA-approved BCL2 inhibitor venetoclax is highly synergistic in AML cell line models with *FLT3* or *KIT* mutations, but also in *FLT3*-mutant AML cells harboring *NRAS*^G12C^. Furthermore, the combination is effective *in vivo*, reducing leukemic burden and improving survival in both cell line-derived xenograft (CDX) and patient-derived xenograft (PDX) AML models. Altogether, these data provide a new mechanistic rationale and preclinical evidence for co-targeting of SHP2 and BCL2 as a therapeutic strategy in *FLT3-* and *KIT-* mutant AML.

## Results

### RMC-4550 is active in *FLT3-* and *KIT-* mutant AML cell lines

As RTK-driven AML relies upon hyperactive RAS/MAPK signaling for survival, we hypothesized that targeting SHP2 would inhibit proliferation of RTK-driven AML cells, as has been previously reported in solid tumors (21, 28–29, 31). To test this hypothesis, we assessed the efficacy of RMC-4550 in nine AML cell lines with RTK or downstream signaling mutations, using a cell viability assay (Fig. 1A). As expected, RMC-4550 inhibited cell proliferation in cell lines with RTK mutations, most potently in the FLT3-ITD mutant cell lines Molm-14 and MV4-11 (IC_50_ 137 nM and 117 nM, respectively) and less potently in the KIT^N822K^ mutant cell lines Kasumi-1 and SKNO-1 (IC_50_ 193 nM and 480 nM, respectively). Pharmacological sensitivity to SHP2 inhibition correlated with the cell lines’ genetic dependency on PTPN11, evaluated in the Cancer Dependency Map portal (DepMap, https://depmap.org) (Fig. 1B) with the exception of U937 cells. Although U937 is a cell line highly dependent on PTPN11, it has an activating *PTPN11*^G60R^ mutation. Substitutions at the G60 residue are shown to bias the open conformation of SHP2 and are associated with resistance to allosteric SHP2 inhibition (34, 35). Accordingly, cell lines with activating mutations in *PTPN11* (U937 – PTPN11^G60R^) and *RAS* (OCI-AML3 – *NRAS*^Q61L^, THP-1 – *NRAS*^G12D^, NOMO-1 – *KRAS*^G13D^, HL-60 – *NRAS*^Q61L^) were resistant to RMC-4550 (IC_50_ > 10 μM, Fig. 1A, Supplementary table 1).

**Figure 1.**
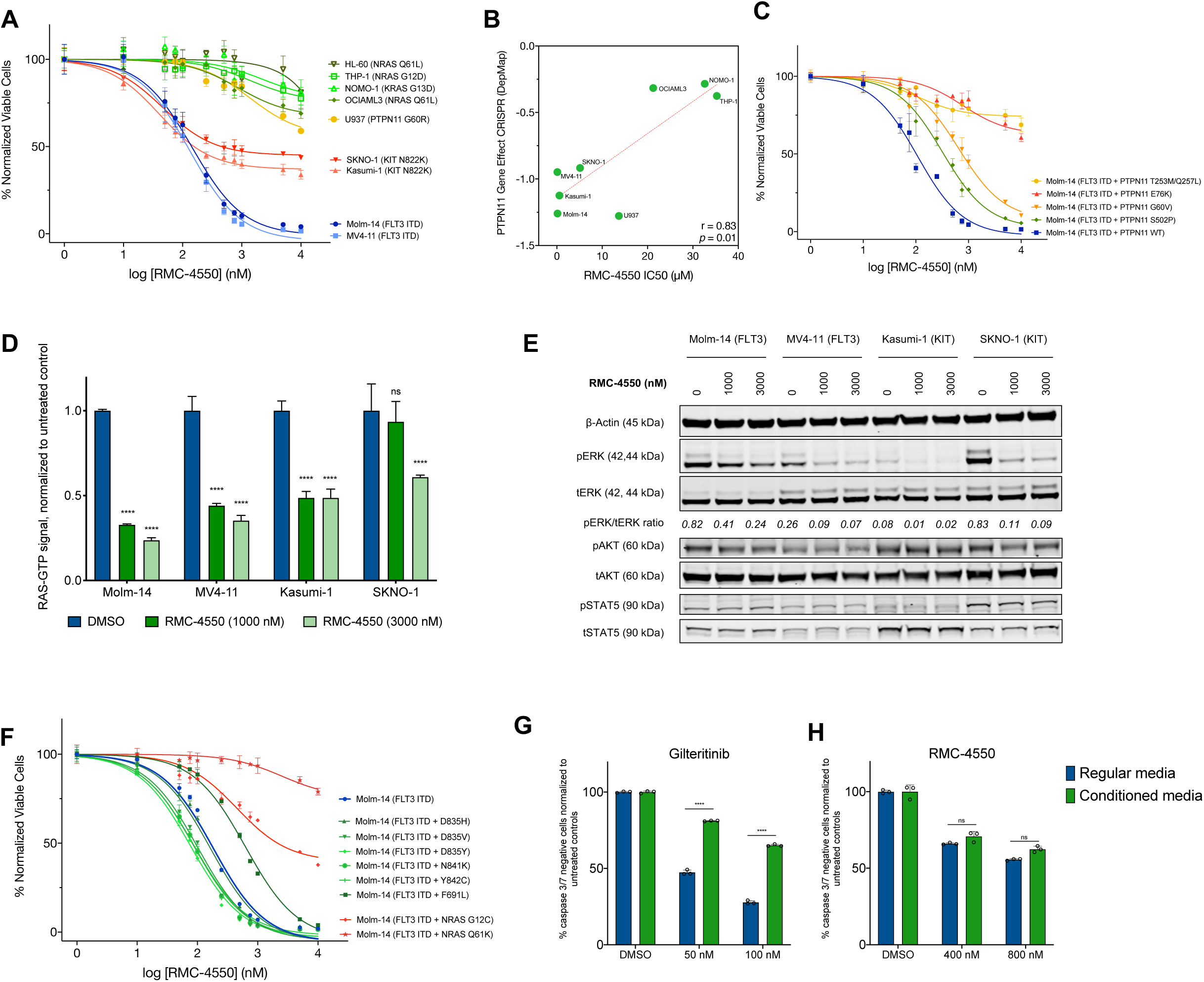
Pharmacologic inhibition of SHP2 has anti-proliferative activity in RTK-driven AML cell lines. **A,** Dose-response curves representing relative proliferation of nine AML cell lines after 48 hours of exposure to serial doses of RMC-4550. Data represent mean ± standard error of mean (SEM) of three technical replicates. **B,** Pearson correlation between sensitivity to RMC-4550 represented by IC_50_ values obtained from a. and genetic dependency on PTPN11 represented as CRISPR gene effect scores from http://depmap.org. Lower scores denote high genetic dependency. Pearson’s r and two-tailed *p* shown **C,** Dose-response curves of Molm-14 cells with doxycycline-inducible expression of PTPN11 variants exposed for 48 hours to serial doses of RMC-4550. Data represent mean ± SEM of three technical replicates. **D,** Colorimetric detection of RAS-GTP levels in four AML cell lines exposed for 90 minutes to RMC-4550. Data represent means of three technical replicates, error bars represent standard deviation (SD); two-tailed ANOVA with Tukey’s correction for multiple comparisons was used for statistical analysis (****, *P* ≤ 0.0001, *ns*, *P* > 0.05). **E,** Western blot analysis of four AML cell lines exposed for 90 minutes to the indicated doses of RMC-4550. Total protein extracts were resolved on a 10% Bis-Tris gel and subjected to immunoblot analysis with the indicated antibodies; actin was used as loading control. For phospho-ERK expression, band intensities from images were normalized to a total ERK control and are shown underneath the relevant bands. **F,** Dose-response curves representing relative proliferation of quizartinib-selected Molm-14 cells with additional mutations in *FLT3* and *NRAS* genes. Data represent mean ± SEM of three technical replicates. **G, H,** Apoptosis measured by flow cytometry in Molm-14 cells cultured in either regular growth media or HS-5 stromal cells-conditioned media after 24 hours of exposure to increasing doses of gilteritinib or RMC-4550. Live, caspase3/7 negative cells were gated and normalized to untreated controls. Data represent means of three technical replicates, error bars represent SD and statistical analysis was performed using unpaired *t* test with Holm-Sidak correction for multiple comparisons (****, *P* ≤ 0.0001; *ns*, *P* > 0.05).

To determine if the anti-proliferative effect of RMC-4550 is due to on-target SHP2 inhibition, we transduced FLT3-ITD+ Molm-14 cells with doxycycline-inducible constructs to overexpress wild-type *PTPN11* or activating *PTPN11* variants bearing activating mutations (35), as well as the double *PTPN11*^T253L/Q257M^ mutation, previously reported to impair the binding of allosteric SHP2 inhibitors (28). The hyperactive PTPN11^E76K^ and the *PTPN11*^T253L/Q257M^ mutants were profoundly resistant to RMC-4550, while the mildly activating *PTPN11*^S502P^ and *PTPN11*^G60V^ mutants exhibited less resistance (Fig. 1C). Furthermore, using a caspase 3/7 assay, we observed that RMC-4550 induced differing degrees of apoptosis in the four parental sensitive lines (Molm-14, MV4-11, Kasumi-1, SKNO-1), but not in the resistant line OCI-AML3 (Supplementary Fig. S1A). At a biochemical level, in RTK-driven sensitive cell lines, RMC-4550 rapidly impaired RAS GTP loading and repressed downstream phosphorylation of ERK (Fig. 1D-E).

### RMC-4550 is active in FLT3 TKI-resistant cell lines

We next evaluated the activity of RMC-4550 in conditions conferring resistance to FLT3 TKIs. RMC-4550 maintained activity against Molm-14 cells expressing FLT3 tyrosine kinase domain (TKD) mutations associated with secondary resistance to FLT3 TKIs (Fig. 1F) (12, 36), including the “gatekeeper” F691L variant and mutations in the FLT3 activation loop. We also evaluated RMC-4550 efficacy in Molm-14 cells harboring secondary *NRAS*^G12C^ or *NRAS*^Q61K^ mutations that emerged during long-term exposure to the FLT3 inhibitor quizartinib. Whereas RMC-4550 retained activity in Molm-14 cells with a *NRAS*^G12C^ mutation, the *NRAS*^Q61K^ variant conferred resistance (Fig. 1F). This finding is in agreement with the previously reported observation that G12C mutants are vulnerable to impaired SOS1/2-mediated exchange induced by SHP2 inhibition (31).

Cell-extrinsic resistance to FLT3i is associated with ERK reactivation induced by bone marrow microenvironment protective cytokines (16). Accordingly, we assessed the ability of RMC-4550 and the FLT3i gilteritinib to induce apoptosis in Molm-14 cells cultured in media conditioned by the immortalized human bone marrow stromal cell line HS5 and in media enriched with FGF2 and FLT3-L at 10 ng/ml (16, 37). When cultured in conditioned and cytokine-enriched media, Molm-14 cells exhibited significantly less apoptosis upon exposure to gilteritinib. In contrast, RMC-4550 was much less vulnerable to resistance to apoptosis induced by conditioned and cytokine-enriched media (Fig. 1F-G, Supplementary Fig. S1B-C). This finding suggests that SHP2 inhibition has the potential to overcome stromal-mediated resistance to FLT3 TKIs.

### SHP2 inhibition alters the transcriptomic profile of RTK-driven AML cell lines

To evaluate the transcriptomic changes induced by SHP2 inhibition and identify potential complementary pathways which could be effectively co-targeted with SHP2i, we performed RNA sequencing (RNA-seq) in two *FLT3*-ITD (Molm-14, MV4-11) and one *KIT*-mutant (SKNO-1) AML cell lines treated with RMC-4550 or vehicle (DMSO). Hierarchical clustering revealed significant changes in global gene expression with SHP2i treatment (Fig. 2A), and principal component analysis (PCA) showed distinct gene expression patterns in samples treated with RMC-4550 compared to vehicle (Supplementary Fig. S2A). Differential gene expression analysis identified 132 overlapping dysregulated genes (fold change > 2, FDR adj. P value < 0.05) in all three cell lines (Fig. 2B). Molm-14 cells exhibited the highest number of differentially expressed genes (3016) and MV4-11 the fewest (464) (Fig. 2B-E). In a cross-cell line pooled analysis, gene set enrichment analysis (GSEA) for hallmark gene signatures (38–39) showed increased expression of genes down-regulated by KRAS activation and decreased expression of MYC targets (Fig. 2F), consistent with expected transcriptomic changes associated with RAS-MAPK signaling inhibition. Cells treated with RMC-4550 also exhibited a marked overexpression of genes implicated in interferon-alpha (IFN-α) and gamma (IFN-*γ*) signaling. Interferons are known to be potent inducers of apoptosis in cancer cells via numerous mechanisms (40–42). Among IFN-stimulated genes (ISGs) upregulated by RMC-4550, IFIT2 (or ISG54; fold change: 3.06 to 18.44) has been shown to induce Bax- and/or Bak-dependent apoptosis via activation of mitochondrial pathway (42). Similarly, RMC-4550 upregulated expression of 2’-5’ oligoadenylate synthetase family members OAS1, OAS2, OAS3, and OASL (fold change: 3.5 to 20). These proteins, under IFN stimulation, induce apoptosis by both degrading intra-cellular RNA via RNAse L (43) and by directly interacting with BCL2 family members (44). Moreover, we observed a significant increase in expression of the IFN-responsive tumor suppressor SAMD9L (54) in all cell lines and of the pro-apoptotic X-linked inhibitor of apoptosis-associated factor 1 (XAF1) in the MV4-11 and SKNO-1 cell lines.

**Figure 2.**
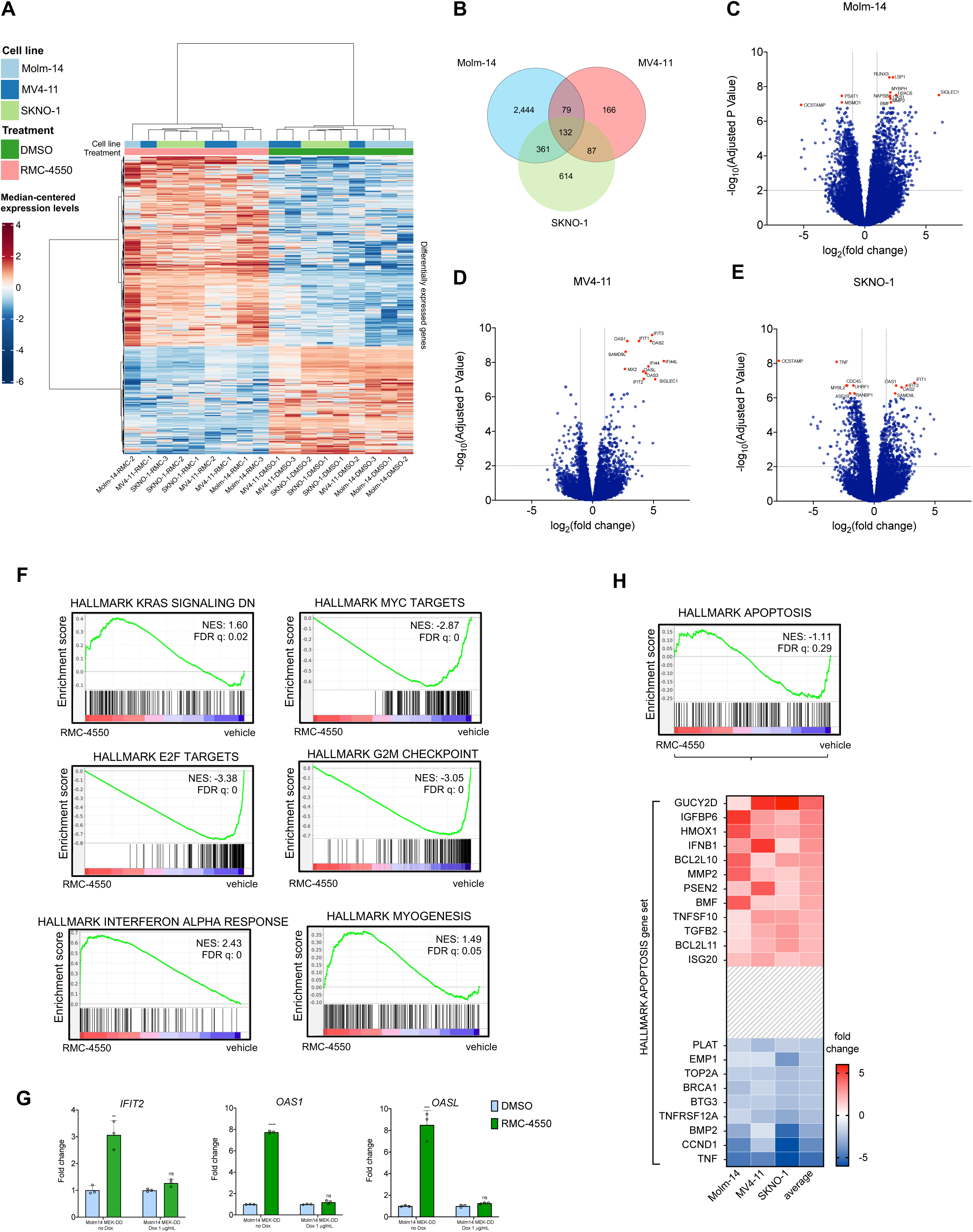
SHP2 inhibition alters the gene expression in FLT3 and KIT mutant AML cell lines. **A,** Heatmap showing gene intensity per sample relative to the average level across all samples. Individual genes are shown on the Y axis and samples are shown on the X axis. Red and blue cells correspond to higher and lower RNAseq levels, respectively. A total of 1,000 genes selected at random from all differentially expressed genes are shown. Samples are n=3 individually treated cell cultures of each cell line: Molm-14, MV4-11 and SKNO-1, treated for 24 hours with either DMSO (vehicle) or RMC-4550 **B,** Venn diagram showing the number of differentially expressed genes in each individual cell line and the overlaps between cell lines. **C, D, E,** Volcano plots displaying differentially expressed genes in each individual cell line. Highlighted genes with above a significance threshold of -log_10_(*p* val) ≥ 7 in Molm-14 and MV4-11 and ≥6 in SKNO-1 cells. **F,** Barcode plots representing GSEA results for the indicated gene sets in all cell lines. Normalized enrichment score (NES) and False Discovery Rates (FDR) were determined by GSEA computational method. **G,** RT-qPCR data representing gene expression fold change relative to DMSO-treated cells of *IFIT2*, *OAS1* and *OASL* genes after 24 hours treatment with RMC-4550. Expression of MEK^DD^ variant was induced with 24 hours treatment with 1 μg/mL of doxycycline. Data represents mean ± SD of three technical replicates; *t* test with Benjamini-Hochberg False Discovery Rate (FDR, Q = 1%) approach was used for statistical significance (****, Q ≤ 0.0001; ***, Q ≤ 0.001; **, *Q* ≤ 0.01; *ns*, Q > 0.01). **H,** Barcode plot of apoptosis gene set and heatmap representing relative expression of genes in this gene set. Genes with an average fold change ≥ 2 or ≤ -2 are shown.

Upregulation of IFN gene sets in response to several kinase inhibitors in RTK-driven cancer cell lines has been previously reported (46). We investigated if the overexpression of IFN targets in our dataset is contingent upon SHP2i ability to suppress RAS/MAPK signaling. RMC-4550 significantly upregulated the gene expression of IFIT-2, OAS-1 and OAS-L in Molm-14 cells, but not in cells with doxycycline-induced overexpression of a dominant active MEK1 allele (MEK^S218D/S222D^, referred to as MEK^DD^) (47) (Fig. 2G), confirming that SHP2i increases transcription of pro-apoptotic IFN targets via suppression of RAS/MAPK signaling.

Consistent with RAS-MAPK inhibition and ISG overexpression, we also noted a significant transcriptional downregulation of genes encoding cell cycle related targets of E2F transcription factors and of genes involved in the G2/M checkpoint, suggesting that SHP2i impairs DNA replication and blocks cell cycle progression (Fig. 2F). An intriguing finding was the enrichment of the myogenesis gene set, a sign of phenotypical remodeling of the contractile cytoskeleton (Fig. 2F). Genetically and pharmacologically interfering with actomyosin contractility has been shown to induce apoptosis in the HL-60 AML cell line and to modulate oncogenic gene expression through the mechano-transducing system of YAP (48). In *EGFR*-mutant NSCLC, hyper-active YAP is associated with resistance to EGFR and MEK1 inhibition and apoptosis evasion through YAP-mediated repression of pro-apoptotic BMF (Bcl2-modifying-factor) (49). However, we did not observe consistent changes in YAP levels with SHP2i treatment in RTK-driven AML cells (Supplementary Fig. S2C). In our dataset, the genes mediating apoptosis were not dysregulated to a significant threshold (FDR = 0.28), likely due to differential expression of genes coding for both pro and anti-apoptotic proteins (Fig. 2H). However, we found the pro-apoptotic BMF and BIM (*BCL2L11*) to be upregulated across all cell lines. Of these, BMF was upregulated to a greater extent on average and was one of the top ten most significantly upregulated genes in Molm-14 cells (fold change: 4.41, FDR: 8.1e-8). *BMF* codes for a BH3-only protein, found at a steady state to be sequestered with actin-associated myosin V motors; upon anoikis (a type of programmed cell death), it detaches from the cytoskeleton, binds and neutralizes BCL2 family members to allow cytochrome C release from mitochondria (50, 51). Anoikis occurs as a consequence of cell detachment from the extracellular matrix (ECM) and cytoskeletal perturbations, preventing ectopic cell growth and re-adhesion (51).

### RMC-4550 increases apoptotic dependency on BCL2 in *FLT3-* and *KIT-* mutant AML

In light of our RNA-seq findings, we sought to confirm transcriptional upregulation of BMF using RT-qPCR. Treatment with RMC-4550 significantly increased the gene expression levels of BMF in a dose-dependent manner in all four investigated cell lines (Molm-14, MV4-11, Kasumi-1, SKNO-1) (Fig. 3A), but not in Molm-14 cells with doxycycline-induced expression of MEK^DD^) (Fig. 3B), highlighting that SHP2i-mediated suppression of MAPK signaling is required for BMF transcriptional upregulation. At the protein level, BMF was increased upon 24 hours exposure to RMC-4550, though the degree of increase varied by cell line. In contrast, consistent changes in other pro and anti-apoptotic proteins were not observed. Upregulation of the pro-apoptotic protein BIM was noted in MV4-11 cells, but not in the other cell lines tested (Fig. 3C). We also noticed decreased levels of the MAPK-dependent anti-apoptotic MCL1 in Molm-14 and SKNO-1 cells (Fig. 3C). In keeping with our RNA sequencing results, Molm14 cells had the greatest degree of SHP2i-induced BMF upregulation on both the transcript and protein level. Consistent with this, Molm-14 cells treated for 24 hours with RMC-4550 exhibited higher cytoplasmic intracellular levels of BMF compared to untreated cells, as shown by immunofluorescent staining (Fig. 3D).

**Figure 3.**
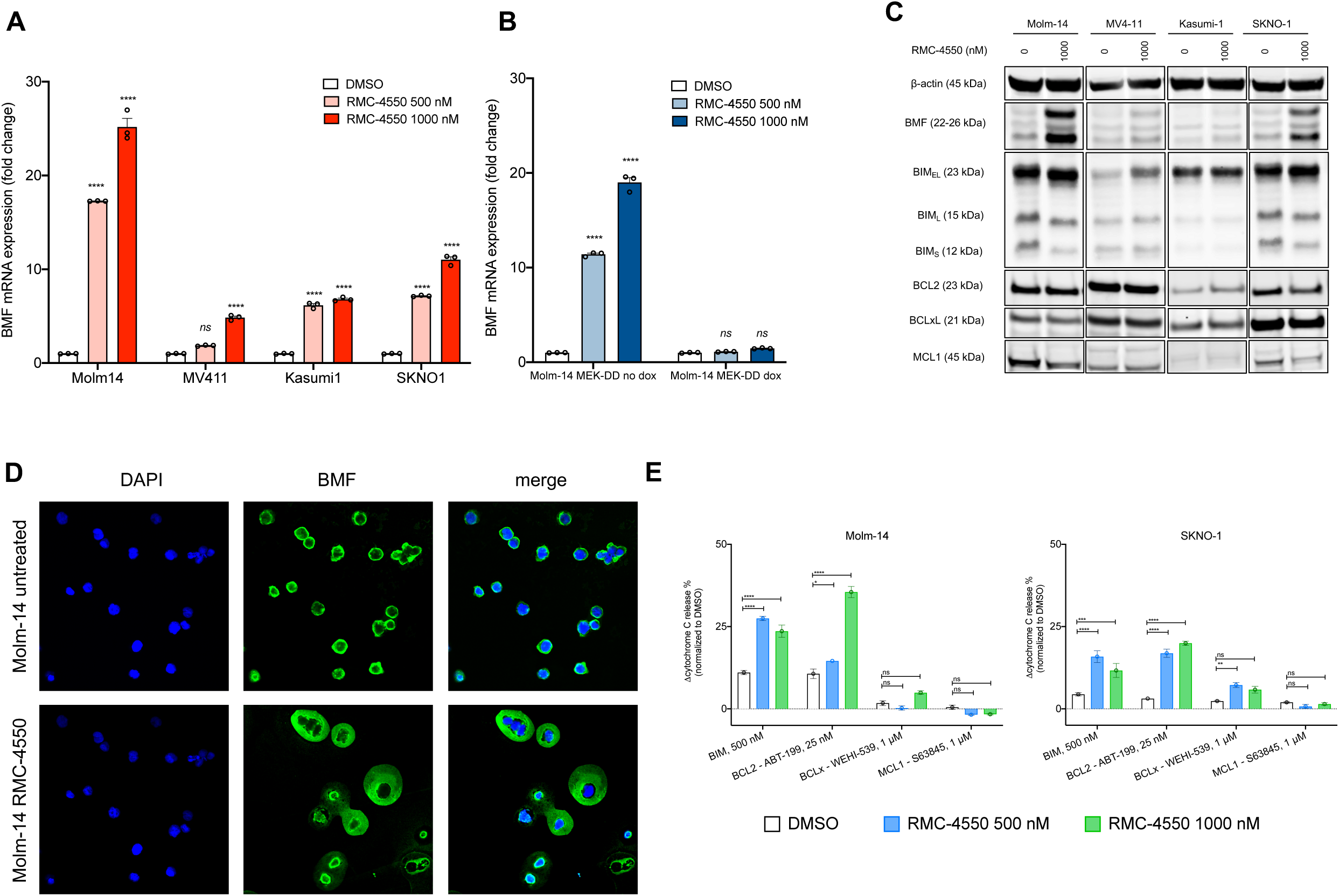
RMC-4550 increases expression of pro-apoptotic BMF and the dependency for apoptosis on BCL2 in RTK-driven AML cell lines. A, RT-qPCR data representing gene expression fold change of BMF gene in Molm-14, MV4-11, Kasumi-1 and SKNO-1 and in B, Molm-14 MEK^DD^ cells after 24 hours of treatment with RMC-4550. Data represent mean ± SD of three technical replicates; two-way ANOVA with Sidak’s correction for multiple comparisons was used for statistical significance C, Western blot analysis of pro- and anti-apoptotic proteins in Molm-14, MV4-11, Kasumi-1 and SKNO-1 exposed to RMC-4550 for 24 hours; actin was used as loading control. D, Immunofluorescence images showing cytoplasmic increase of BMF (green) after 24 hours of treatment with RMC-4550 in Molm-14 cells. DAPI (blue) was used as a nuclear stain. Images were acquired at a 60X magnification using confocal microscopy. E, iBH3 profiling data representing normalized cytochrome C release following 16 hours of treatment with DMSO or RMC-4550 and exposure to either BIM peptide or BCL2, BCLxL and MCL1 inhibitors in Molm-14 and SKNO-1 cells; Data represent mean ± SD of three technical replicates; two-way ANOVA with Sidak’s correction for multiple comparisons was used for statistical significance (****, *P* ≤ 0.0001, ***, *P* ≤ 0.001, **, *P* ≤ 0.01, * *P* ≤ 0.05, *ns*, *P* > 0.05).

BMF exerts its pro-death response via binding BCL2 family members. Amid its binding partners, BMF has the highest affinity for BCL2 (52). In order to assess the apoptotic dependency to individual BCL2 family proteins, we used a BH3 profiling assay (53) to quantify the mitochondrial outer membrane permeabilization (MOMP) as a result of exposure to either BH3 peptides (BIM, 0.5 μM) or inhibitors (BCL2 – ABT-199/venetoclax, 25 nM; BCLxL – WEHI539, 1 μM; MCL1 – S63845, 1 μM) in Molm-14 and SKNO-1 cells. Treatment with RMC-4550 increased global apoptotic priming (measured by increased cytochrome C release after exposure to BIM peptide) and apoptotic dependency on BCL2 (measured by increased cytochrome C release after exposure to a low dose of ABT-199 (25 nM)) in both cell lines. These findings suggest that upon SHP2 inhibition, the apoptotic dependency of these AML cells shifts towards BCL2, implying co-inhibition of SHP2 and BCL2 may be a mechanistically complementary approach to induce apoptosis in RTK-driven AML.

### Pharmacologic co-targeting of SHP2 and BCL2 has synergistic anti-proliferative activity in *FLT3*- and *KIT*-mutant AML

Having pinpointed BCL2 dependency as a vulnerability of RTK-driven AML cells treated with SHP2i, we hypothesized that RMC-4550 may sensitize these cells to pharmacological inhibition of BCL2 with venetoclax. To test this hypothesis we treated Molm-14, Molm-14 *NRAS*^G12C^, MV4-11, Kasumi-1 and SKNO-1 cells with both agents and performed formal assessment of drug synergy using a cell viability readout (CellTiterGlo) and the SynergyFinder 3.0 computational tool (54). The combination was highly synergistic in all investigated cell lines (Fig. 4A). We observed the highest synergy scores in Molm-14 *NRAS*^G12C^ and SKNO-1 (Bliss scores of 31 and 42 respectively), cells with the lowest sensitivity to RMC-4550 as a single agent (Fig. 1A, 1F). Furthermore, the combination of RMC-4550 and venetoclax induced a striking increase of apoptosis in comparison to individual single agents (Fig. 4B, S4A). RMC-4550 and venetoclax were also more synergistic in Molm-14 cells than gilteritinib and venetoclax (Supplementary Fig. S3B), an active combination that achieved high modified composite complete remission (mCRc) rates in a phase Ib clinical investigation in patients with R/R AML (55).

**Figure 4.**
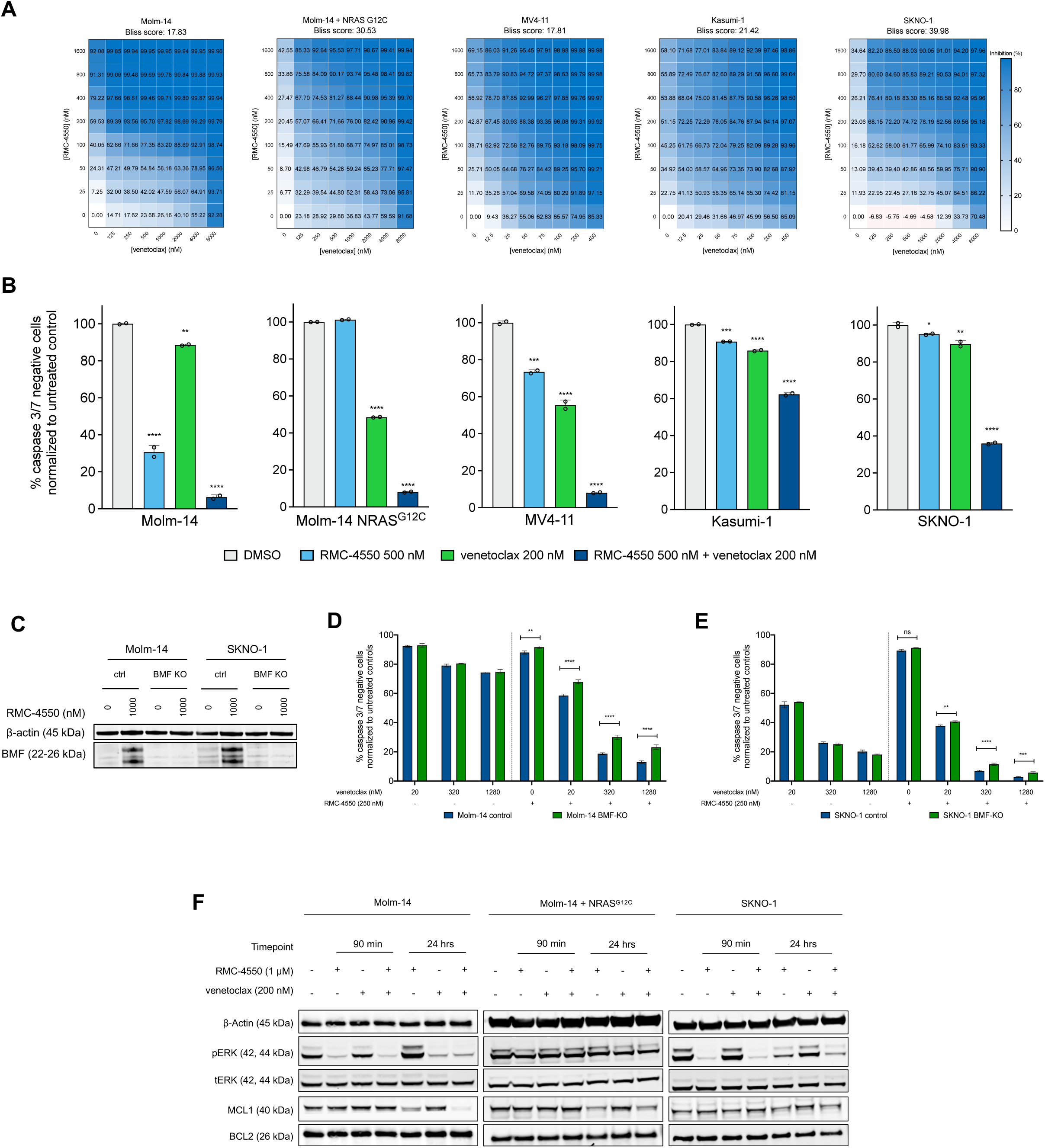
Co-targeting SHP2 and BCL2 is a synergistic therapeutic approach in *FLT3*- and *KIT*-mutant AML. **A**, Dose-response matrices representing normalized cell viability inhibition following 48 hours of treatment with increasing doses of RMC-4550 and venetoclax in the indicated cell lines. Synergy scores were computed using Bliss method within Synergy Finder v3.0 software. **B,** Apoptosis measured by flow cytometry after 24 hours for treatment with RMC-4550 and venetoclax at indicated concentrations. Results depict live, caspase3/7 negative cells normalized to untreated controls. Data represent mean ± SD of two technical replicates; statistical analysis was performed using one-way ANOVA test with Dunnett correction for multiple comparisons. **C,** Western blot analysis of BMF protein after 24 hours of treatment with RMC-4550 in Molm-14 and SKNO-1 cells with BMF CRISPR knock-out; actin was used as loading control. **D,E,** Apoptosis measured after 24 hours of treatment with RMC-4550 and venetoclax in Molm-14 and SKNO-1 BMF KO cells compared to control cells. Data represent mean ± SD of three technical replicates; two-way ANOVA with Sidak’s correction for multiple comparisons was used for statistical analysis. **F,** Western blot analysis showing phoshpo-ERK and MCL1 dysregulations after treatment for 90 minutes and 24 hours with RMC-4550 and venetoclax; actin was used as loading control. (****, *P* ≤ 0.0001, ***, *P* ≤ 0.001, **, *P* ≤ 0.01, * *P* ≤ 0.05, *ns*, *P* > 0.05).

Having demonstrated that SHP2 inhibition upregulates BMF, a potent BCL2 repressor, we aimed to investigate whether BMF is required for the synergistic activity of RMC-4550 and venetoclax. Using CRISPR-Cas9 ribonucleoprotein transfection, we knocked out *BMF* in Molm-14 and SKNO-1 cells. Western blotting confirmed the ablation of BMF expression in knock-out (KO) cells upon SHP2 inhibition (Fig. 4C). We treated cells with intact *BMF* and *BMF* KO with either 250 nM of RMC-4550 or DMSO, then exposed them for 24 hours to increasing doses of venetoclax and measured apoptosis. Compared to DMSO, treatment with RMC-4550 increased apoptosis in response to venetoclax in all cell lines, but to a lesser extent in BMF KO than in BMF competent cells (Fig. 4D-E). This difference underlines that competent BMF is required for maximal sensitization to BCL2 inhibition by RMC-4550 in both RTK-driven AML models.

Although RMC-4550 initially suppressed ERK phosphorylation in both Molm-14 and SKNO-1 cell lines (Fig. 1C), we note rebound activation of pERK after 24 hours of treatment (Fig. 4F). Notably, the addition of venetoclax successfully reversed the rebound compared to RMC-4550 alone in both cell lines and downregulated MCL1 further in Molm-14 cells (Fig. 4F). These observations are in line with similar findings reported in Molm-14 cells co-treated with gilteritinib and venetoclax (56). Moreover, we found that RMC-4550 and venetoclax mildly decreased pERK and downregulated MCL1 even in the presence of the *NRAS*^G12C^ mutation (Fig. 4F).

To investigate whether the synergic induction of cell death is a consequence of MAPK signaling suppression by SHP2i, we treated doxycycline-inducible *MEK*^DD^ overexpressing Molm-14 cells with RMC-4550 or vehicle, then exposed them to venetoclax and measured apoptosis. Overexpression of dominant-active MEK^DD^ rescued cells from cell death induced by the SHP2i-BCL2i combination (Supplementary Fig. S3C). In contrast, doxycycline-inducible overexpression of a dominant-active AKT variant, *AKT*^E17K^, did not prevent apoptosis induction in Molm-14 cells treated with the combination (Supplementary Fig. S3D). These findings highlight that effective inhibition of RAS-MAPK, but not AKT, signaling is necessary for SHP2i-mediated sensitization to venetoclax. In support of this observation, we found that MEK inhibitor trametinib and venetoclax were similarly synergistic in inhibiting cell proliferation of all cell lines investigated (Supplementary Fig. S3E), in agreement with previous data reporting pre-clinical synergy between the MEK inhibitor cobimetinib and venetoclax in AML (57).

### The combination of RMC-4550 and venetoclax is active in cell line-derived xenograft (CDX) models of AML

We next sought to determine whether the synergistic activity of RMC-4550 and venetoclax observed *in vitro* could be recapitulated *in vivo* using CDX models. To emulate multiple distinct clinical scenarios, we evaluated AML cells with *FLT3*-ITD – Molm-14, Molm-14 cells with acquired *NRAS*^G12C^ mutation, and *KIT*-mutant – Kasumi-1 cells. Cells were transfected with a firefly-luciferase-containing lentiviral construct, then injected into NOD/SCID/IL2Rγnull (NSG) female mice. Upon engraftment, mice were randomized in four groups receiving treatment with either vehicle, RMC-4550 (30 mg/kg, orally, once daily, p.o., q.d.), venetoclax (100 mg/kg, p.o, q.d.) or their combination, 5 days per week, for a total of 28 days (Fig. 5A). Using in-vivo bioluminescence imaging (BLI), we observed that combination treatment reduced *in vivo* tumor burden (Fig. 5B, 5D, Supplementary Fig. 4A-D). In all three CDX models, mice in the combination arm had significantly enhanced survival compared to the control group and to monotherapies (Fig. 5C, 5E-F). Collectively, these data demonstrate *in vivo* combinatorial efficacy of SHP2 and BCL2 inhibition in *FLT3-* and *KIT-* mutant AML models, as well as in a model of AML resistant to FLT3i due to secondary *NRAS*^G12C^ mutation.

**Figure 5.**
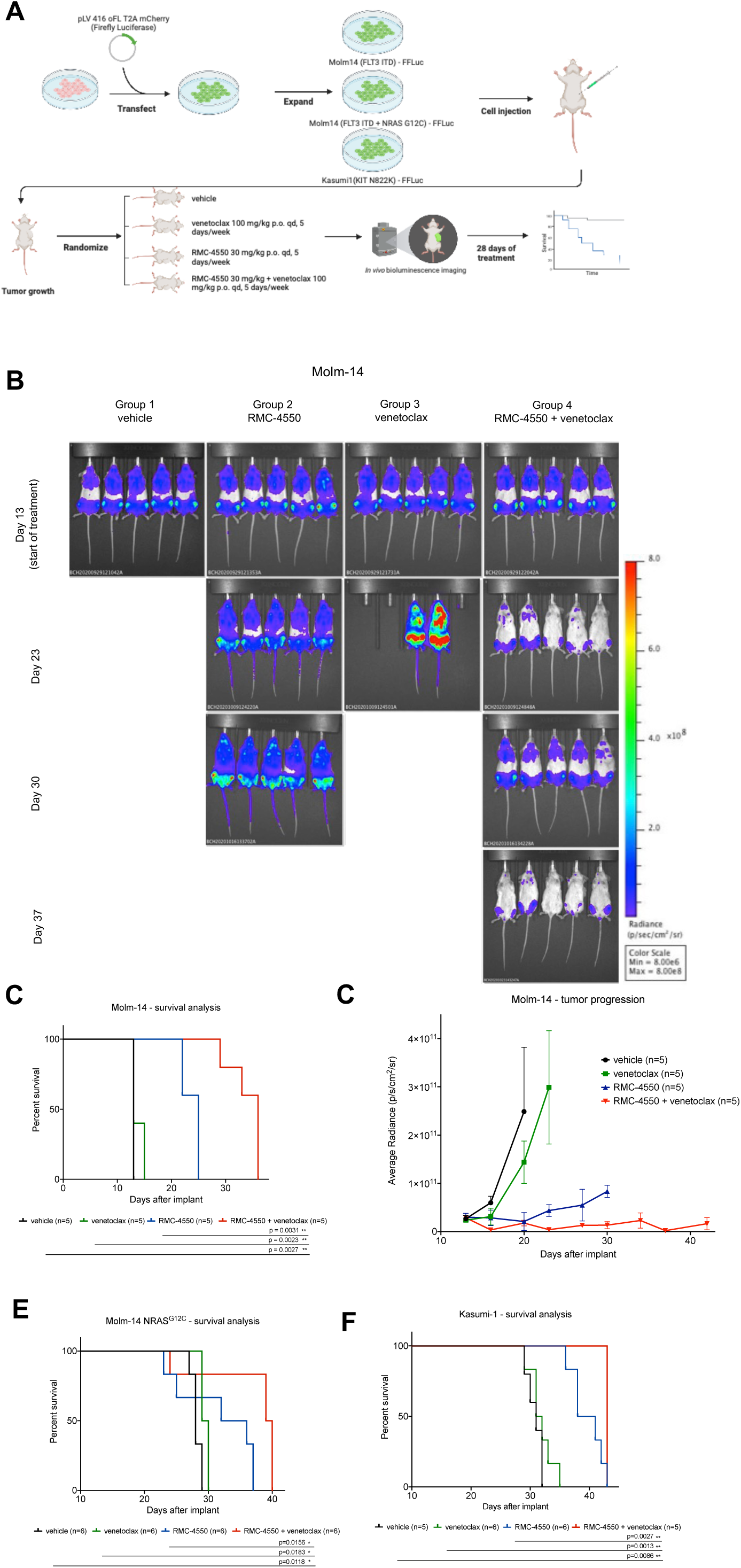
Combination therapy with RMC-4550 and venetoclax is effective *in vivo* in CDX AML models. **A,** Schematic representation of the CDX *in vivo* study design. Image created with BioRender.com. **B,** Representative images of *in vivo* bioluminescence imaging (BLI) assessment of NSG mice engrafted with luciferase-tagged Molm-14 cells over the course of the trial. **C,E,F,** Kaplan-Meier curves representing survival data of subjects from each treatment group in **C,** Molm-14, **E,** Molm-14 *NRAS*^G12C^, **F,** Kasumi-1 CDX studies; log-rank (Mantel-Cox) test was used for comparison of survival curves. **D,** Quantification of BLI data from the Molm-14 CDX study, data represents mean ± SD (n=5/group).

### RMC-4550 and venetoclax combination is effective in patient-derived AML xenografts (PDX) and primary samples

We next aimed to verify the anti-leukemic activity of RMC-4550 and venetoclax in AML patient-derived xenograft (PDX) models. To determine a tolerable dosing schedule, we initially treated non-tumor bearing NSG-SGM3 (NSG mice with transgenic expression of human GM-CSF, IL-3, and SCF in an NSG background) mice with two doses (20 mg/kg and 30 mg/kg) of RMC-4550 in combination with venetoclax (100 mg/kg). At the end of 28 days of treatment, no significant body weight changes or treatment-related toxicities were reported with either dosing regimen (Supplementary Fig. S5A), and the 30 mg/kg dosing strategy was selected for further PDX studies.

We next tested the combination in two PDX models established from patients with *FLT3-* ITD AML (clinical genotyping data in Table S1). AML cells were injected into busulfan-conditioned NSG-SGM3 mice, as previously reported (58). After engraftment, mice were randomized into treatment groups: vehicle, RMC-4550 (30 mg/kg, p.o., q.d), venetoclax (100 mg/kg, p.o., q.d.) and the combination of the two agents (Fig. 6A). Flow-cytometric quantification of human CD45 (hCD45)-expressing cells in murine peripheral blood obtained from intermediate timepoint bleedings was used to monitor disease progression. The combination therapy reduced the percentage of circulating hCD45+ cells compared to both vehicle control and single-agent venetoclax in both PDX models and versus single agent RMC-4550 in the PDX#1 model (Fig. 6B, Supplementary Fig. S5B). After 28 days of treatment or at pre-determined humane study endpoint, mice were sacrificed and hCD45+ were enumerated in murine cardiac blood, spleen, and bone marrow by flow cytometry. Combination therapy led to significant suppression of leukemic burden in comparison to vehicle and venetoclax in all assessed organs (Fig. 6C-D). The activity of RMC-4550 alone varied by assessed tissue, but the combination demonstrated significant suppression of hCD45+ cells compared to RMC-4550 alone, in spleen for one xenograft model (Fig. 6C) and bone marrow for the other (Fig. 6D). RMC-4550 alone or in combination also reduced spleen weight and length (Fig. 6E, Supplementary Fig. S5C-D). Immunohistochemical staining of murine spleen samples at trial termination additionally showed a marked decreased of hCD45+ cells in mice receiving the combination (Fig. 6F).

**Figure 6.**
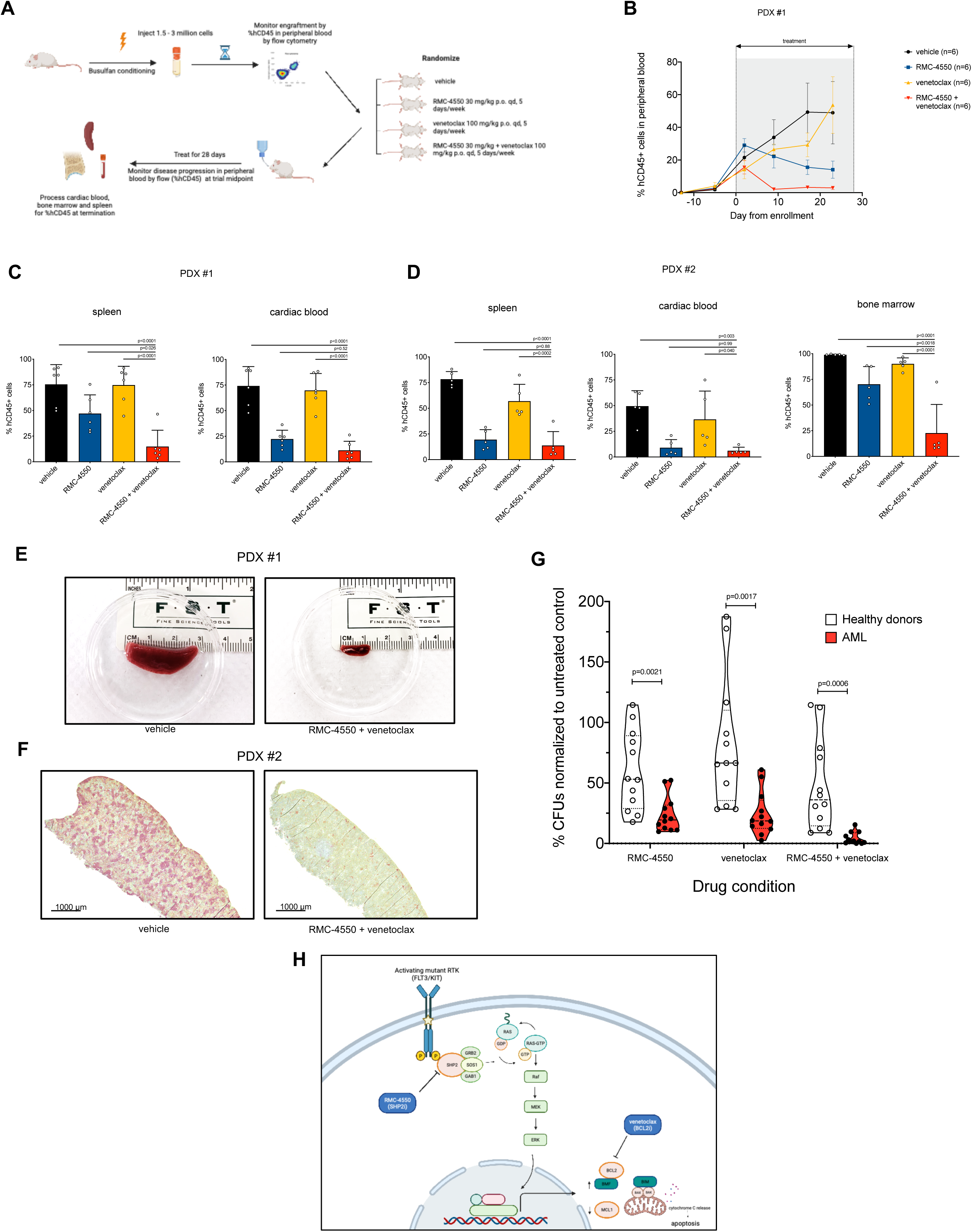
Simultaneous SHP2 and BCL2 inhibition is effective in *FLT3*-mutant AML PDX models and in primary AML samples. **A,** Schematic representation of the PDX *in vivo* study design. Image created with BioRender.com **B,** quantification of hCD45+ cells over the course of treatment in the four treatment groups in the PDX #1 study. **C, D,** Quantification of hCD45+ in target organs at the study termination (PDX #1 - n=6/group, PDX #2 – n=5/group). Data represent mean ± SD; one-way ANOVA with Tukey correction for multiple comparisons was used for statistical analysis. **E,** Representative images of spleens harvested from subjects in the vehicle or RMC-4550 + venetoclax treatment groups in PDX #1 study. **F,** Representative images of immunohistochemical staining for hCD45 in spleen samples from subjects in the vehicle or RMC-4550 + venetoclax treatment groups in PDX #2 study. **G,** Violin plots showing differences in colony-forming unit (CFU) normalized counts in primary samples derived from healthy donors (n=4) or AML patients (n=4) treated for 14-16 days with RMC-4550, venetoclax or their combination. Data represents three technical replicates for each sample; unpaired *t* test with Holm-Sidak correction for multiple comparisons was used for statistical analysis. **H,** Proposed mechanism for the synergistic cell-death inducing activity of simultaneous SHP2 and BCL2 inhibition. Image created with BioRender.com

Finally, in an effort to determine if the combination of RMC-4550 and venetoclax would be expected to suppress normal hematopoiesis, we evaluated the colony-forming unit (CFU) potential of primary bone marrow mononuclear cells (BMMC) harvested from four healthy donors compared to four FLT3 mutant AML patients in the presence of DMSO, RMC-4550 (500 nM), venetoclax (200 nM), or both inhibitors. Whereas the combination dramatically inhibited CFU formation in AML samples, healthy donor samples retained CFU formation, suggesting potential for a therapeutic index (Fig. 6G).

## Discussion

Acquired resistance to FLT3 inhibitors is a major challenge of the current therapeutic paradigm of *FLT3*-mutant AML. Single-cell genomic analyses of patients treated with FLT3 inhibitors have illustrated the heterogenous nature of on-target and off-target resistance mechanisms (12–13, 15). Our group and others have demonstrated that under FLT3 inhibition, clones that re-activate the RAS/MAPK pathway account for clinical resistance in a large proportion of patients (13, 15, 59). Pre-clinical studies suggest that feedback activation of ERK signaling limits FLT3i efficacy (60) and that microenvironment factors protecting from FLT3i also converge on reactivation of downstream MAPK and PI3K/AKT signaling (16–17). Collectively, these data imply that RAS/MAPK signaling is a primary mediator of survival signaling in *FLT3*-mutant AML.

SHP2 inhibition decreases oncogenic RAS-RAF-MEK-ERK signaling through disruption of SOS1-mediated RAS-GTP loading (21). The addition of SHP2i to MEK inhibitors successfully reversed feedback activation of RTK leading to resistance to various in solid malignancies (32, 61–62). In *FLT3*-mutant AML, SHP2i showed promising single agent activity in cell lines and murine models (28, 30); however, SHP2i have not been yet clinically investigated in patients with AML. Furthermore, single agent targeting of the MAPK pathway in AML has been largely ineffective clinically (63–65). In this study, we demonstrate that targeting SHP2 exposes a vulnerability of RTK-driven AML by increasing apoptotic priming and BCL2 dependency through multiple MAPK-dependent mechanisms, resulting in increased sensitivity to venetoclax-induced cell death (Fig. 6H). Our *in vivo* and *in vitro* findings thus provide robust pre-clinical evidence that coupling SHP2 blockade with the clinically available BCL2i venetoclax is a synergistic, effective approach against *FLT3*- and *KIT*-mutant AML, and can overcome several mechanisms of adaptive resistance to FLT3i.

We demonstrate that targeting SHP2 simultaneously addresses multiple heterogenous FLT3i resistance mechanisms, including secondary *FLT3* TKD mutations, exchange-sensitive RAS mutations and AML-protective microenvironment crosstalk signals associated with early resistance. We notably show that RMC-4550 retained activity against TKI-resistant Molm-14 cells with *NRAS*^G12C^, in agreement with published data suggesting that oncogenic *RAS* G12C variants are sensitive to upstream modulation of SOS1-mediated nucleotide exchange (21, 31). Moreover, we show that RMC-4550 successfully overcame the paracrine-mediated resistance to FLT3i-induced apoptosis conferred by microenvironmental soluble factors, such as stromal secreted cytokines, or exogenous FLT3-L and FGF2 (16).

Using RNA-seq, we identified a pro-death transcriptomic signature induced by SHP2 inhibition, consisting of upregulation of targets under the negative control of ERK, suppression of MYC targets, consistent with the findings of another group (30), and repression of E2F targets. Additionally, we noticed a significant upregulation of IFN-*α−* and IFN-*γ*-stimulated genes. Interferons are known to be potent inducers of apoptosis in cancer cells via numerous mechanisms (41–42). Upregulation of IFN target genes following MAPK inhibition has been previously reported in oncogenic signaling cancers, but was associated with a drug-induced immune reprogramming resulting in a senescence-like state and immune evasion (46). While the function of SHP2 inhibition in AML-extracellular matrix adhesion remains to be explored, we observed significant upregulations of the myogenesis gene set and the pro-apoptotic BMF, implicating a role for anoikis as a potential event triggering apoptosis. In the context of anoikis, BMF is known for translocating from the contractile cytoskeleton to the cytoplasm transducing a mechanical signal to the apoptotic machinery. However, we observed a transcriptional upregulation of BMF that was reversed in the setting of MEK activation, suggesting it relies upon effective ERK suppression. High intracellular amounts of BMF lead to neutralization of anti-apoptotic proteins, particularly BCL2, facilitating BIM/BAX/BAK-mediated apoptosis (50, 52). Simultaneous with BMF overexpression, we noted downregulation of MCL1. Using BH3 profiling assays we demonstrated that SHP2 inhibition increases overall sensitivity to BIM-induced apoptosis, but also the dependency of leukemia cells on BCL2. Collectively, these data suggest that SHP2i uncovers an unexpected susceptibility of RTK-driven AML, rendering AML cells dependent on BCL2 for cell death and therefore increasing their sensitivity to venetoclax. Indeed, the combination of RMC-4550 and venetoclax synergized in inhibiting leukemia proliferation, inducing apoptosis and preventing ERK reactivation. Of note, the combination dramatically increased caspase levels even in AML cells where either agent only modestly induced apoptosis. Recently, it was reported that clinical resistance to venetoclax in AML is associated with monocytic differentiation and a low *BCL2*/*MCL1* gene expression ratio, whereas MEKi could sensitize monocytic AML cells to venetoclax (66–67). We show that RMC-4550 inhibits MAPK signaling and downregulates MCL1, suggesting a mechanism whereby SHP2i can potentially enhance the pharmacologic response to BCL2i in venetoclax-resistant AML.

A recent study methodically investigating the mechanism of synergy between gilteritinib and venetoclax in Molm-14 cells, attributed the drug synergy exclusively to BIM activation (56). The proposed mechanism is that venetoclax displaces BCL2 from BIM, while gilteritinib downregulates MCL1, allowing thereby-available, free BIM to initiate apoptosis. In addition to MCL1 downregulation, we observed that BMF is a contributing factor to the synergy between RMC-4550 and venetoclax, as Molm-14 BMF knock-out cells showed less apoptosis than BMF competent cells when treated with the combination. We found that BMF overexpression in response to SHP2 inhibition relies entirely on the ability of RMC-4550 to suppress MAPK signaling, considering that expression of BMF is unchanged in SHP2i-resistant MEK^DD^ cells. Importantly, the combination of RMC-4550 and venetoclax was tolerable and effective in inhibiting leukemia progression and improving survival in clinically-relevant CDX and PDX AML models.

Of note, RMC-4550 and venetoclax did not elicit a synergic induction of apoptosis in the setting of MEK^DD^ overexpression, emphasizing the necessity of efficient MAPK signaling suppression for cytotoxicity. While upregulation of interferon signaling, BMF and downregulation of MCL1 appear to be partially responsible for induction of apoptosis and BCL2 dependency, the full molecular mechanisms of RMC-4550 and venetoclax synergy remain to be elucidated. Nonetheless, these data suggest that other strategies co-targeting RAS/MAPK signaling and BCL2 would be similarly synergistic. Our study supports the idea that RAS/MAPK signaling is a critical mediator of survival signaling in RTK-driven AML. SHP2 inhibition suppress this pathway to induce pro-apoptotic changes and an increase in BCL2 dependency that may be opportunistically exploited with concurrent BCL2 inhibition. Our findings provide strong mechanistic rationale to support clinical investigation of a SHP2 and BCL2 inhibitor combination in RTK-driven AML.

## Methods

### Cell lines

Molm-14, MV4-11, OCIAML-3, U937, HL-60, NOMO-1, THP-1 were a gift from Dr. Scott Kogan (University of California, San Francisco, San Francisco, CA). Kasumi-1 and SKNO-1 cell lines were purchased from ATCC. HS5 cells were a gift from Dr. Neil Shah (University of California, San Francisco, San Francisco, CA). Molm-14 *NRAS*-mutant and FLT3 TKD–mutant cells were generated by culturing Molm-14 parental cells *in vitro* in media containing escalating doses of quizartinib (0.5–20 nmol/L, ref. 68). Quizartinib-resistant cells were subcloned and Sanger sequencing was performed for mutation confirmation. To generate *PTPN11-*mutant and MEK^DD^ Molm-14 doxycycline-inducible expression cell lines, PTPN11 and MEK variants were cloned into a Gateway tetracycline-inducible destination lentiviral vector, pCW57.1 (Addgene). The cloning was performed by Twist Biosciences. Lenti-X 293T cells (Takara Bio) were infected using Lipofectamine 3000 (Invitrogen) and cultured for 48 hours in DMEM with 10% FBS. The supernatant was harvested, concentrated using Lenti-X concentrator (Takara Bio), then used to infect cell lines. Forty-eight hours following lentiviral infection, cells were selected with puromycin. To generate luciferase-tagged cell lines we transfected parental Molm-14, quizartinib-resistant Molm-14 *NRAS^G12C^*, and Kasumi-1 cells with the Firefly-luciferase expressing plasmid pLV 416 oFL T2A mCherry, a gift from Neil Shah (University of California, San Francisco, San Francisco, CA).

Cell lines were cultured in RPMI-1640 (Gibco) with 10% fetal bovine serum (FBS) and 1% penicillin/streptomycin/L-glutamine (Gibco), with the exception of Kasumi-1 and HL-60 cells, cultured with 20% FBS. Culture media for SKNO-1 cell line was supplemented with human GM-CSF (10 ng/mL, PeproTech). Cells tested negative for *Mycoplasma* by the MycoAlert PLUS Mycoplasma Detection Kit (Lonza). Experiments were performed within three months of cell line thawing. Authentication of all cell lines was performed at the University of California, Berkeley, DNA Sequencing Facility, using short tandem repeat DNA profiling.

### Patient samples

Coded primary AML samples were collected, stored, and studies performed after approval of research protocols from the University of California San Francisco Hematologic Malignancies Tissue Bank or the Children’s Hospital of Philadelphia Center for Childhood Cancer Research biorepository by institutional review boards. Informed, written consent according to the Declaration of Helsinki was obtained from patients prior to tissue collection.

### Compounds

RMC-4550 was provided by Revolution Medicines. Venetoclax (S8048), gilteritinib (S7754), WEHI-539 (S7100) and trametinib (S2673) were purchased from Selleckchem. S63845 (HY-100741) was purchased from MedChem Express. BIM peptide was purchased from New England Peptide.

### Cell viability assays

For cell viability experiments, cells were seeded in three technical replicates in 96-well plates and exposed to increasing concentrations of indicated drug for 48 hours. For doxycycline-inducible cell line experiments, cells were were stimulated for 24 hours with doxycycline 1 mg/mL and maintained in the same concentration of doxycycline for the duration of experiments. Cell viability was assessed using CellTiter-Glo Luminescent Cell Viability Assay (Promega), luminescence was measured on a Molecular Devices iD3 multimode plate reader and normalized to untreated controls.

### Apoptosis assays

Cells were cultured in appropriate culture media, then seeded in three technical replicates in 96-well clear plates and exposed to increasing concentrations of indicated drug for 24 hours. CellEvent Caspase-3/7 Green Flow Cytometry Kit (Thermo Fisher) was used to measure fraction of apoptotic cells by flow cytometry (BD LSR Fortessa with HTS sampler). Percentage of caspase-3/7 negative cells in each treatment condition was normalized to vehicle-treated control populations (FlowJo software v9, BD Biosciences).

### Conditioned media experiments

To generate conditioned media, 10^6^ HS5 cells were plated in 10 mL of complete RPMI in a 10-cm dish and media harvested after 72 hours, cleared by centrifugation and filtration through a 0.22-mm membrane. Molm-14 cells were cultured in conditioned media, complete RPMI and RPMI supplemented with FGF-2 and FLT3-L (10 ng/mL) each, as previously described (16, 37), then exposed to indicated doses of inhibitors for 24 hours and assessed for apoptosis.

### Active RAS detection assay

For RAS-GTP detection, a Ras GTP-ase immunosorbent assay (G-LISA) Activation Assay Kit (#BK131, Cytoskeleton) was used, as per manufacturer’s protocol. In brief, 1 x 10^6^ cells were incubated with drug treatments for 90 minutes, then cell lysates were harvested. 25 μg of total protein per condition were incubated in triplicate in wells coated with Raf1-RBD. Wells were then washed, and incubated with an anti-Ras antibody (#GL-11) and subsequently with a HRP-conjugated secondary antibody (#GL-02). Absorbance readout at 490 nM was used to assess the colorimetric response.

### Immunoblotting

Cells were plated in appropriate media and treated with the indicated concentrations of inhibitors. After 90 minutes or 24-hours incubation, cells were washed in PBS and lysed in buffer (50 mmol/L HEPES, pH 7.4, 10% glycerol, 150 mmol/L NaCl, 1% Triton X-100, 1 mmol/L EDTA, 1 mmol/L EGTA, and 1.5 mmol/L MgCl_2_) supplemented with protease and phosphatase inhibitors (EMD Millipore). The lysates were clarified by centrifugation, quantitated by BCA assay (Thermo Scientific) and normalized. 20 mg of protein were loaded on 10% Bis-Tris gels then transferred to nitrocellulose membranes. Immunoblotting was performed using the following antibodies from Cell Signaling Technology: anti-β-Actin (8H10D10, #3700), anti-Erk1/2 (3A7, #9107), anti-phoshpo-Erk1/2 (Thr202/Tyr204, #9101), anti-Akt (#9272), anti-phospho Akt (Thr308, #9275), anti-Stat5 (D2O6Y, #94205), anti-phospho-Stat5 (Tyr 694, #9351), anti-Bcl2 (124, #15071), anti-Bcl-xL (#2762), anti-Mcl-1 (#4572), anti-Bim (C34C5, #2933), anti-BMF (E5U2J, #50542). Nitrocellulose membranes were subsequently incubated in a solution of secondary antibodies (IRDye 800CW Goat anti-Rabbit, #935-32211, IRDye 680RD Goat anti-Mouse, #935-68070, LI-COR Biosciences) and scanned on an Odyssey CLx infrared imaging system (LI-COR Biosciences). When needed, band intensities were quantified using the Image Studio software.

### RT-qPCR

Cells were incubated for 24 hours with indicated drug concentrations and total RNA was isolated using RNeasy Mini kit (QIAGEN). cDNA was synthesized from 500 ng total RNA using SuperScript III reverse transcriptase (Invitrogen) and qPCR was performed in 384-well plates using Taqman Gene Expression assays (GAPDH: Hs02758991_g1, OASL: Hs00984387_m1, OAS1: Hs00973637_m1, IFIT2: Hs01922738_s1, BMF: Hs00372937_m1,) on a QuantStudio 6 instrument (Thermo Fisher). Differential gene expression was calculated using the 2^−ΔΔCt^ method with GAPDH as a housekeeping control gene in three technical replicates.

### RNA-seq

Molm-14, MV4-11 and SKNO-1 cell were exposed to drug treatments, in three biological replicates per condition. RNA samples with RIN scores between 8.4 and 10 were used to generate libraries and sequenced by BGI Genomics using DNBseq platform. mRNA was purified using oligo(dT)-attached magnetic beads and reverse transcribed. The cDNA fragments were end-repaired, 3’-adenylated, and PCR-purified after adaptor ligation. PCR products were quantified on 2100 Analyzer instrument (Agilent), heat-denatured and circularized by a splint oligo sequence to generate circular DNA nanoball (DNB) libraries. DNBs were loaded into a patterned nanoarray and sequenced by Probe-Anchor Synthesis (cPAS), generating 100 bp paired-end reads. Quality metrics for raw sequencing data were evaluated with FastQC. Reads were aligned to the human reference genome GRCh38 and gene counts were quantified using STAR (69). Raw counts were normalized using the trimmed mean of M-values (TMM) method. Differential gene expression analysis was carried out using limma-voom (70) and the threshold for differential gene expression was an adjusted *P* < 0.05, corrected for multiple comparisons using the false discovery rate adjustment (FDR) and a fold-change in expression ≥ 2. R version 4.0.4 running on Ubuntu 20.04 LTS was used for data processing and analysis. Gene set enrichment analysis (GSEA v4.2.3, ref. 38) was used to identify enriched functional gene sets.

### Immunofluorescence

Molm-14 cells were treated with either vehicle solution or 1 μM of RMC-4550. 5e^4^ cells were cyto-spun on glass slides using a Cytospin 3 (Shandon) cytocentrifuge, then fixed with 4% paraformaldehyde, then permeabilized and blocked with a solution of 0.1% Triton-X and 10% normal goat serum (#50197Z, Thermo Fisher) in PBS. The slides were incubated overnight with an anti-BMF antibody solution (#NBP1-76658, Novus Biologicals) 10 μg/mL in PBS with 0.01% Tween-20 and 1% BSA, washed, then incubated with a secondary antibody (#A-21244, Invitrogen). Coverslips were mounted using ProLong Gold Antifade Mountant with DAPI (#P36935, Thermo Fisher). Images were acquired with on a Nikon A1R-MP confocal microscope with a 60x magnification objective and processed using Fiji (ImageJ v 2.3.0).

### BH3 profiling

The BH3 profiling of AML cell lines was carried out as previously described (53). In brief, cells were treated for 24 hours with the indicated doses of RMC-4550, then incubated in 96-well plates in a solution containing 0.002% digitonin and either Bim peptide or Bcl2, BclxL or Mcl1 inhibitors, fixed and stained with an anti-cytochrome C (clone 6H2.B4, # 612310, BioLegend). FACS acquisition was performed on a BD Fortessa LSR instrument with HTS sampler. Cytochrome C release was measured by gating cytochrome C negative populations, normalized to vehicle-treated control populations (FlowJo software v9, BD Biosciences).

### CRISPR Cas9 KO

BMF knock-out was performed using a Gene Knockout Kit v2 (Synthego), using a multi-guide sgRNA approach. Ribonucleoprotein (RNP) was assembled by combining a 180 pmol mixture of three sgRNA guides (Sequences G1: 5’-AAGGCCAGGGCCACAGCAGU-3’, G2: 5’-AAGCUCCCGGGUUGGGUCAC-3’, G3: 5’-GGAGCCAUCUCAGUGUGUGG-3’) with 30 pmol of *Cas9-2NLS* nuclease from *S. pyogenes* in Opti-MEM (Gibco) media and transfected into Molm-14 and SKNO-1 cell lines using a MaxCyte ATX system. RNP-transfected cells and cells transfected with Cas9 only, used as negative controls, were downstream cultured in complete media. Knock-out efficiency was determined by Inference of CRISPR Edits (ICE) analysis (71) and functionally assessed by Western Blot.

### Cell line-derived xenografts

All *in vivo* xenograft animal studies were performed in full accordance with UCSF Institutional Animal Care and Use Committee (IACUC) at the UCSF Preclinical Therapeutics Core (PTC) or the Children’s Hospital of Philadelphia IACUC and Laboratory Animal Facility. NOD/SCID/IL2Rγnull female mice were injected with cell line-derived xenografts consisting of 5 x 10^6^ Firefly-luciferase-tagged Molm-14, Molm-14 *NRAS^G12C^* and Kasumi-1 cells suspended in PBS. Tumor volume progression was assessed bi-weekly using a Xenogen IVIS Spectrum imaging system (Perkin Elmer) after injection of 150 mg/kg of D-luciferin (GoldBio). Upon luminescence-detectable engraftment, animals were randomized into four groups (n=5 or n=6): a control group receiving vehicle, a group receiving RMC-4550, 30 mg/kg, a group receiving venetoclax 100 mg/kg, and a group receiving the combination of the two agents. Treatments were administered by oral gavage, daily, 5 times a week during weekdays, for 28 days. Animals were sacrificed in compliance to IACUC protocols at the study termination or at humane endpoint, as per UCSF Laboratory Animal Resource Center (LARC) veterinary advice.

### Patient-derived xenograft models

Patient derived xenograft models of human *FLT3*-ITD AML were generated in NSG or triple transgenic NOD.scid.Il2Rγc*null*-SGM3 (NSGS) mice (Jackson Laboratory) and serially passaged. Mice were conditioned with busulfan (Sigma) 25mg/kg intraperitoneally on days -2 and -1. Mice were injected with 1 to 1.5 million cells after 1 day of rest. Mice were monitored for weight change weekly during entirety of the trial. Mice were bled by submandibular puncture weekly starting on week 4 from injection and analyzed by FACS for human (hCD45) and mouse CD45 (mCD45). Mice were randomly assigned to a treatment arm once they reached greater than or equal to (0.5% hCD45+ cells / total events) by FACS. Treatment arms were defined as following: vehicle (oral gavage, 5X per week), RMC-4550 (30 mg/kg, oral gavage, 5X per week), venetoclax (100 mg/kg, oral gavage, 5X per week), combination: RMC-4550 (30 mg/kg) and venetoclax (100 mg/kg, oral gavage 5X per week). Mice were treated for 28 days (20 days of dosing), during which they were monitored daily for disease progression and every other week by FACS. Mice were euthanized after 28 days or when they reached humane endpoint, whichever was sooner. Mice were processed upon endpoint and cardiac blood, bone marrow and spleen were analyzed by FACS for hCD45/mCD45. Spleen fragments were processed for immunohistochemical staining.

### Immunohistochemistry (IHC) Staining

IHC staining was performed at the UCSF Histology and Biomarkers Core on a Ventana BenchMark Ultra instrument using Discovery reagents (Ventana Medical Systems) according to manufacturer’s instructions, except as noted. After deparaffinization and antigen-retrieval, slides were incubated with primary antibody anti-CD45 (1:200, clone: D9M8I, Product ID: 13917 from Cell Signaling). The primary antibody was detected with Discovery Purple kit. Finally, slides were counterstained with hematoxylin (Thermo Scientific Shandon Instant Hematoxylin cat. 6765015). Images were acquired using a Zeiss Axio Scanner.Z1 slide scanner with a 20X Plan-Apochromat objective and processed using Zeiss Zen Lite v3.6.

### Colony-forming unit assay

Primary samples from leukapheresis products harvested from either healthy individuals or patients with AML were thawed in DMEM media (Gibco) supplemented with 20% FBS, 2mM EDTA and 500 μg DNase I. 5 x 10^4^ to 2.5 x 10^5^ cells were incubated with DMSO, RMC-4550 (500 nM), venetoclax (200 nM) or the combination of both drugs in methylcellulose media enriched with human cytokines (recombinant human SCF, GM-CSF, G-CSF, IL-3, IL-6, Epo, #HSC005, R&D Systems). The methylcellulose mixture was plated in three replicates to 35 mm culture dishes using regular syringes with 16 gauge non-stick needles and incubated at 37°C, 5% CO_2_ for 14-16 days. Colony forming units were counted using an Olympus CKX53 light microscope with 4X magnification objective and the counts were normalized to untreated controls.

### Statistical analysis

Details on statistical analysis of experiments can be found in the figure legends. Unpaired *t* tests were used to compare independent groups. One-way or two-way analysis of variance (ANOVA) tests with were used to compare more than two groups, and correction for multiple comparisons was applied when needed. Statistical significance is denoted as follows: *, P ≤ 0.05; **, P ≤ 0.01; ***, P ≤ 0.001; ****, P ≤ 0.0001, unless stated otherwise. Data are reported as mean ± SD, unless otherwise specified. For cell viability readouts, dose-response curves and absolute IC_50_ values were calculated using a three parameters nonlinear regression model. Drug synergy assessment was performed using the Bliss method within SynergyFinder v3.0 (https://synergyfinder.fimm.fi/, ref. 54), with LL4 curve fitting and cell viability readout data. For *in vivo* studies, no statistical methods were used to predetermine sample sizes, but our sample sizes are similar to those previously published (29, 30). Data collection and analysis were not performed blind to the experimental conditions. GraphPad Prism software (v8.3.1) was used for statistical analysis and data plotting. BioRender.com was used for to draw graphic illustrations.

## Data availability

Raw and processed RNA-seq data have been deposited at the National Center for Biotechnology Information Gene Expression Omnibus under the accession number <u>GSE217359</u>.

## Authors’ contributions

**B. Popescu:** conceptualization, data curation, formal analysis, investigation, methodology, software, validation, visualization, writing – original draft, writing – review & editing; **C. Stahlhut:** resources, project administration, writing – review & editing; **T. C. Tarver:** formal analysis, investigation, methodology; **S. Wishner:** investigation, validation, visualization; **B. J. Lee:** resources, project administration; **C. A. C. Peretz:** resources, methodology, writing – review & editing; **C. Luck:** software, formal analysis, writing – review & editing; **P. Phojanakong:** investigation, methodology; **J. A. Camara Serrano:** investigation, visualization; **H. Hongo:** data curation, investigation; **J. M. Rivera:** investigation, methodology; **S. Xirenayi:** data curation, investigation; **J. A. Chukinas:** resources; **V. Steri:** resources, investigation, methodology, supervision; **S. K. Tasian:** resources, writing – review & editing; **E. Stieglitz:** methodology, resources, supervision, writing – review & editing; **C. C. Smith:** conceptualization, data curation, funding acquisition, methodology, project administration, resources, writing – original draft, writing – review & editing;

## Ackowledgements

The authors thank the Smith lab members for scientific input and technical support throughout the project; Kevin Shannon for critical reading of this manuscript; Fios Genomics for assistance with bioinformatic analysis of RNA-seq data; Noura Ismail for technical support with confocal microscopy imaging; Christopher Chien and Nirali Shah at the National Cancer Institute (NCI) for assistance with AML specimens for xenograft studies. This work was supported in part by Revolution Medicines. C. A. C. Peretz was funded by a Young Investigator Award from the Alex’s Lemonade Stand Foundation. C. Luck was supported by the NIGMS Predoctoral Training in Biomedical Sciences T32 grant (C.L., T32 GM 136547). The Preclinical Therapeutic Core (PTC) at UCSF (P. Phojanakong, J. A. Camara Serrano, V. Steri) is supported by Cancer Center Support Grant (CCSG - NCI/NIH), award P30CA082103. S. K. Tasian is a Scholar of the Leukemia & Lymphoma Society and holds the Joshua Kahan Endowed Chair in Pediatric Leukemia at the Children’s Hospital of Philadelphia. C. C. Smith is a Leukemia & Lymphoma Society Scholar in Clinical Research and a Damon Runyon-Richard Lumsden Foundation Clinical Investigator supported (in part) by the Damon Runyon Cancer Research Foundation (CI-99–18).

