## Supplementary Data for "Allosteric SHP2 Inhibition Increases Apoptotic Dependency on BCL2 and Synergizes with Venetoclax in *FLT3-* and *KIT-* Mutant AML"

Supplementary Figure S1

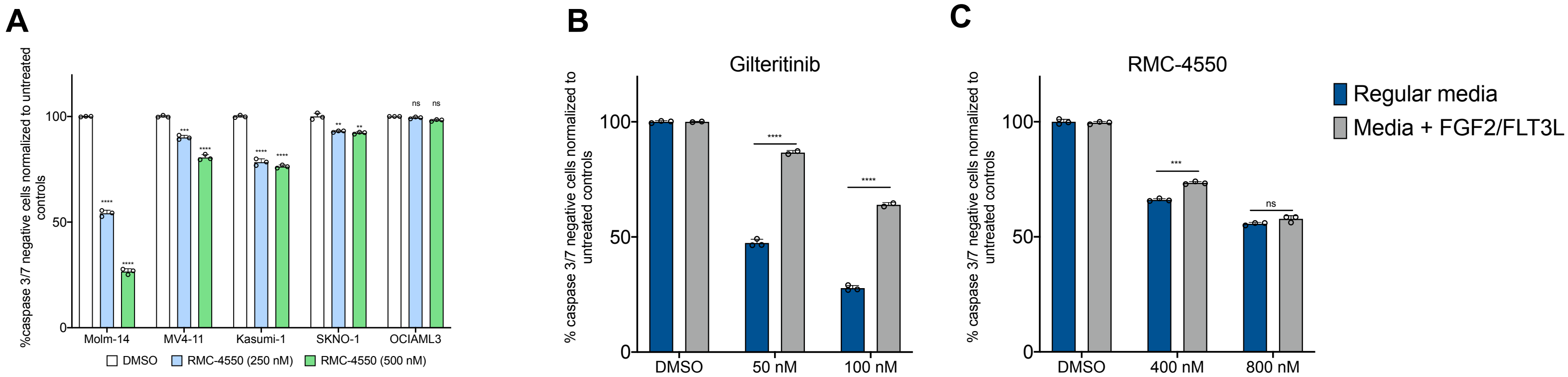

**Supplementary figure S1. SHP2 inhibition has anti-leukemic activity in RTK-driven AML cell lines. A,** Apoptosis assay measuring live, caspase 3/7 negative cells in AML cell lines after 24 hours of treatment with indicated doses of RMC-4550. Statistical analysis was performed using unpaired *t* test with with Holm-Sidak correction for multiple comparisons. **B,** Apoptosis assay after 24 hours of treatment with indicated doses of gilteritinib and RMC-4550 in Molm-14 cells cultured in either regular growth medium or medium supplemented with 10 ng/mL FGF2 and 10 ng/mL FLT3-L. Statistical analysis was performed using two-way ANOVA with Sidak's correction for multiple comparisons. Data represented as mean ± SD of three technical replicates. (****, $P \leq 0.0001$, ***, $P \leq 0.001$, **, $P \leq 0.01$, * $P \leq 0.05$, *ns*, $P > 0.05$).

Supplementary Figure S2

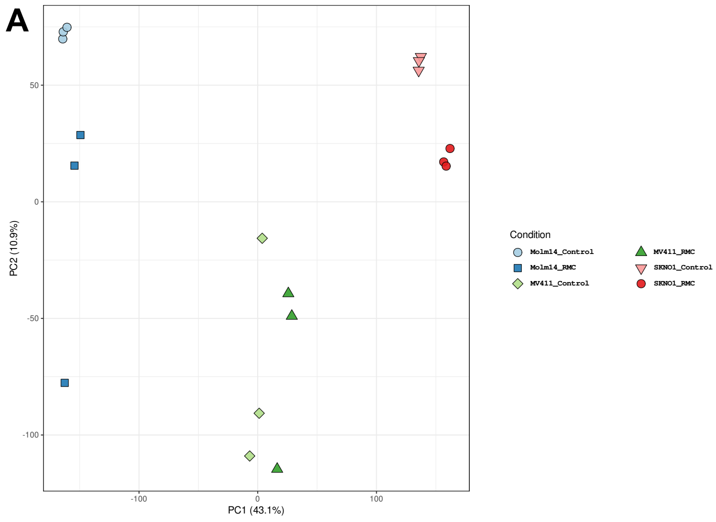


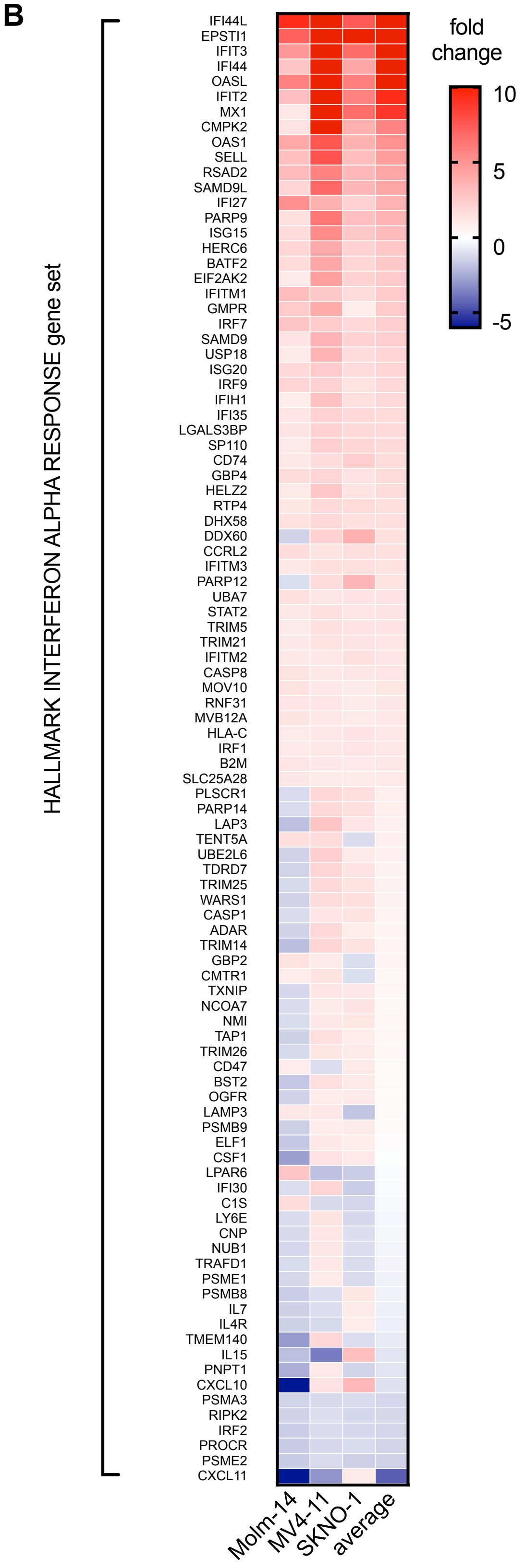


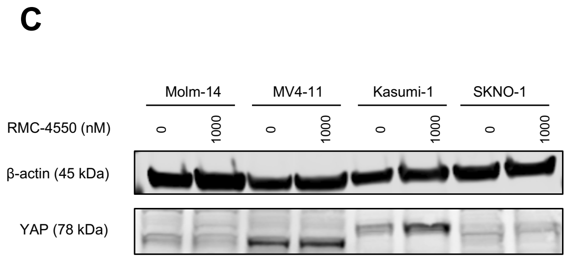

**Supplementary figure S2. SHP2 inhibition alters the transcriptomic profile of RTK-driven AML. A,** Principal component analysis (PCA) scatter plot for the first two principal components from the normalized reads data. Each symbol corresponds to individually treated cell cultures of each cell line (n=3), treated with either DMSO or RMC-4550. **B,** heatmap representing relative expression of genes in the Hallmark Interferon Alpha Response (MSigDB) gene set in all three cell lines. **C,** Western blot analysis of YAP protein expression in four cell lines; actin was used as loading control.

### Supplementary Figure S3

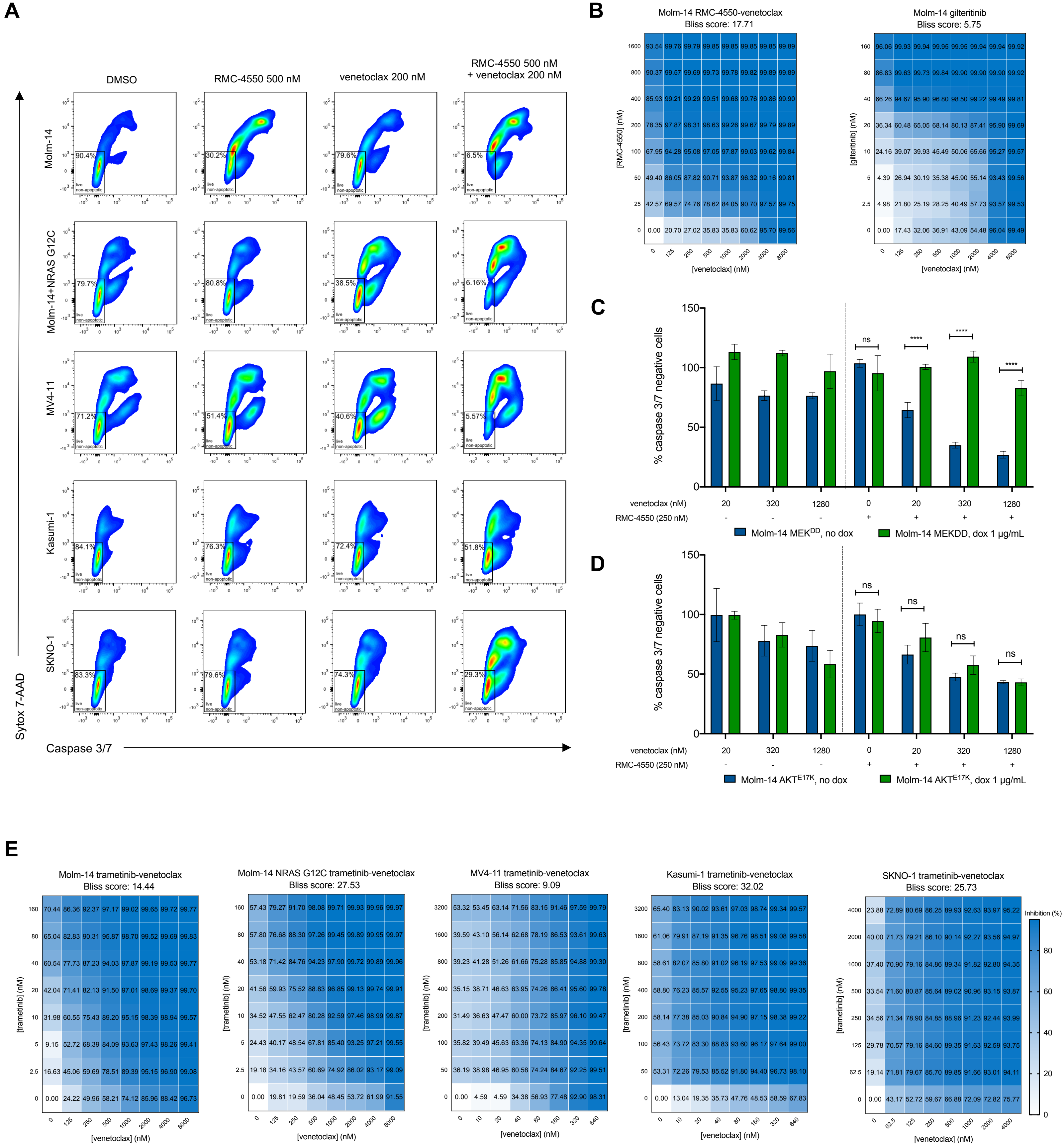

**Supplementary figure S3. Co-targeting SHP2 and BCL2 is a synergistic therapeutic approach in *FLT3-* and *KIT-* mutant AML. A**, Apoptosis assay of cell lines after 24 hours of treatment with RMC-4550, venetoclax and combination; representative gating is shown. **B, E,** Dose-response matrices representing normalized cell viability inhibition following 48 hours of treatment with increasing doses of gilteritinib and venetoclax **(b)** and trametinib and venetoclax **(E)** in the indicated cell lines. Synergy scores were computed using Bliss method within Synergy Finder v3.0 software. **C, D,** Apoptosis measured after 24 hours of treatment with RMC-4550 and venetoclax in Molm-14 cells overexpressing $MEK^{DD}$ **(E)** or $AKT^{E17K}$ **(D)** compared to control cells. Data represents mean ± SD of three technical replicates; two-way ANOVA with Sidak's correction for multiple comparisons was used for statistical analysis (****, $P \leq 0.0001$; ***, $P \leq 0.001$; **, $P \leq 0.01$; *, $P \leq 0.05$; *ns*, $P > 0.05$).

Supplementary Figure S4

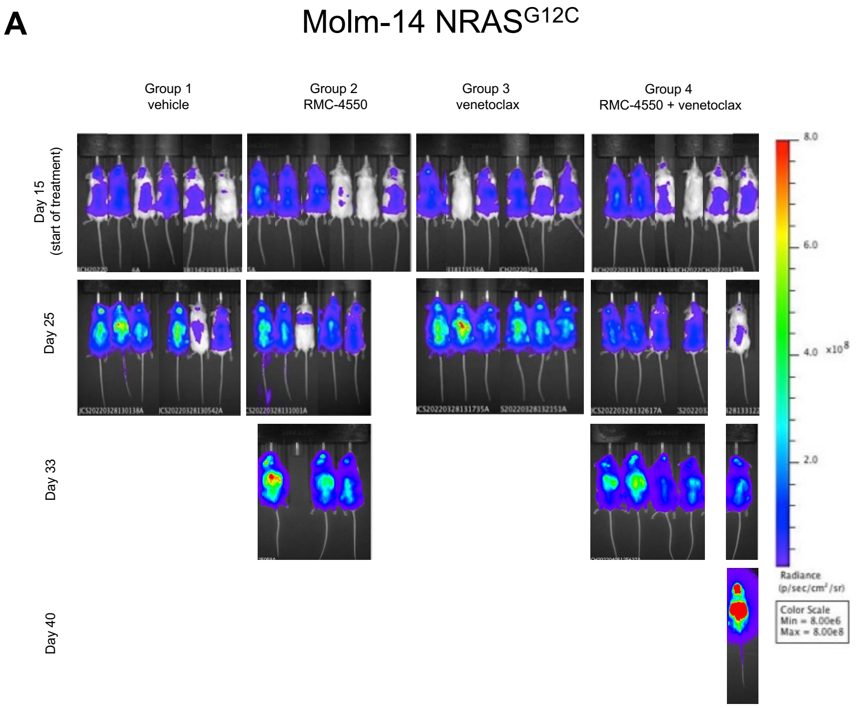


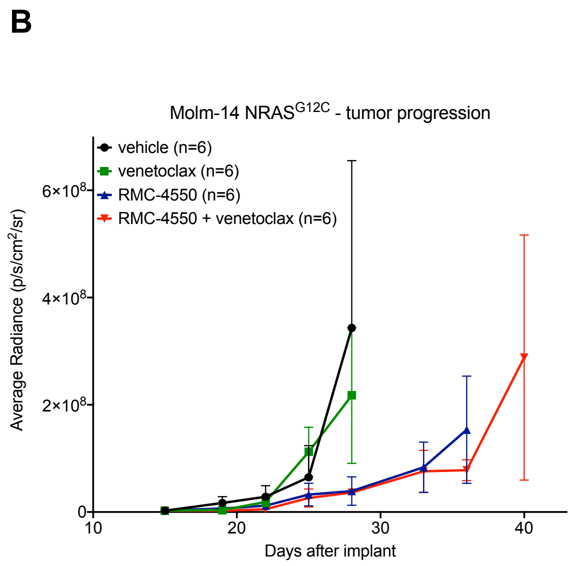


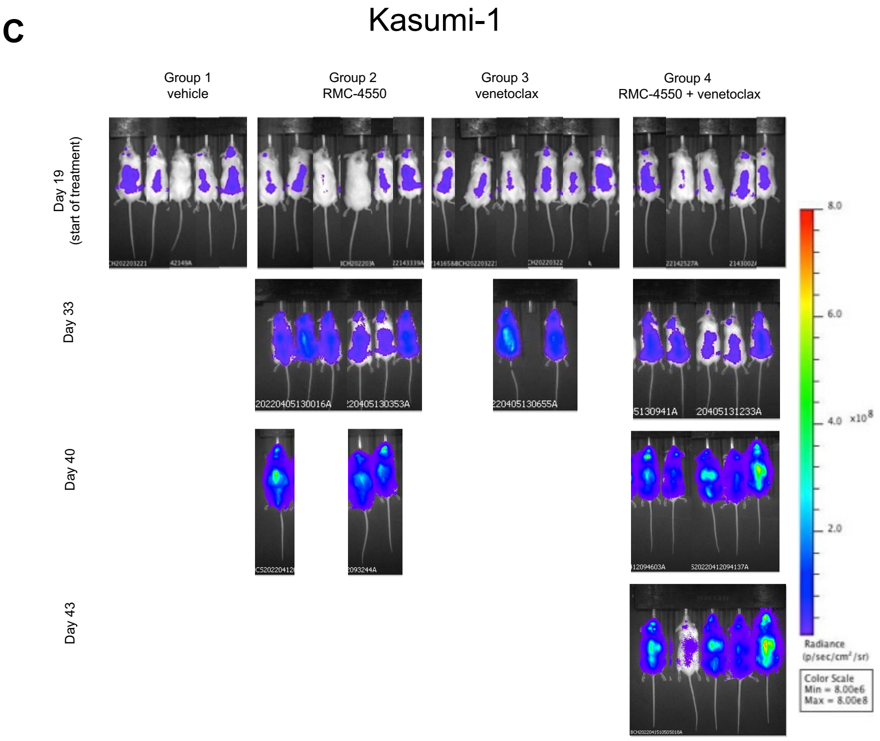


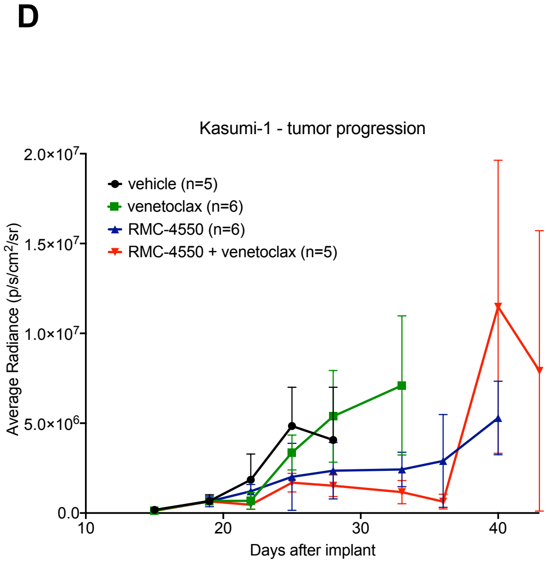

**Supplementary figure S4. Combination therapy with RMC-4550 and venetoclax is effective *in vivo* in CDX AML models. A, C,** Representative images of *in vivo* bioluminescence imaging (BLI) assessment of NSG mice engrafted with luciferase-tagged Molm-14 *NRAS*$^{G12C}$ (n=6/group, **A**) and Kasumi-1 (n=5/group, **C**) cells over the course of the trial. **B, D,** Quantification of BLI data from the Molm-14 *NRAS*$^{G12C}$ (**B**) and Kasumi-1 (**D**) CDX studies, data represents mean ± SD (n=5/group).

Supplementary Figure S5

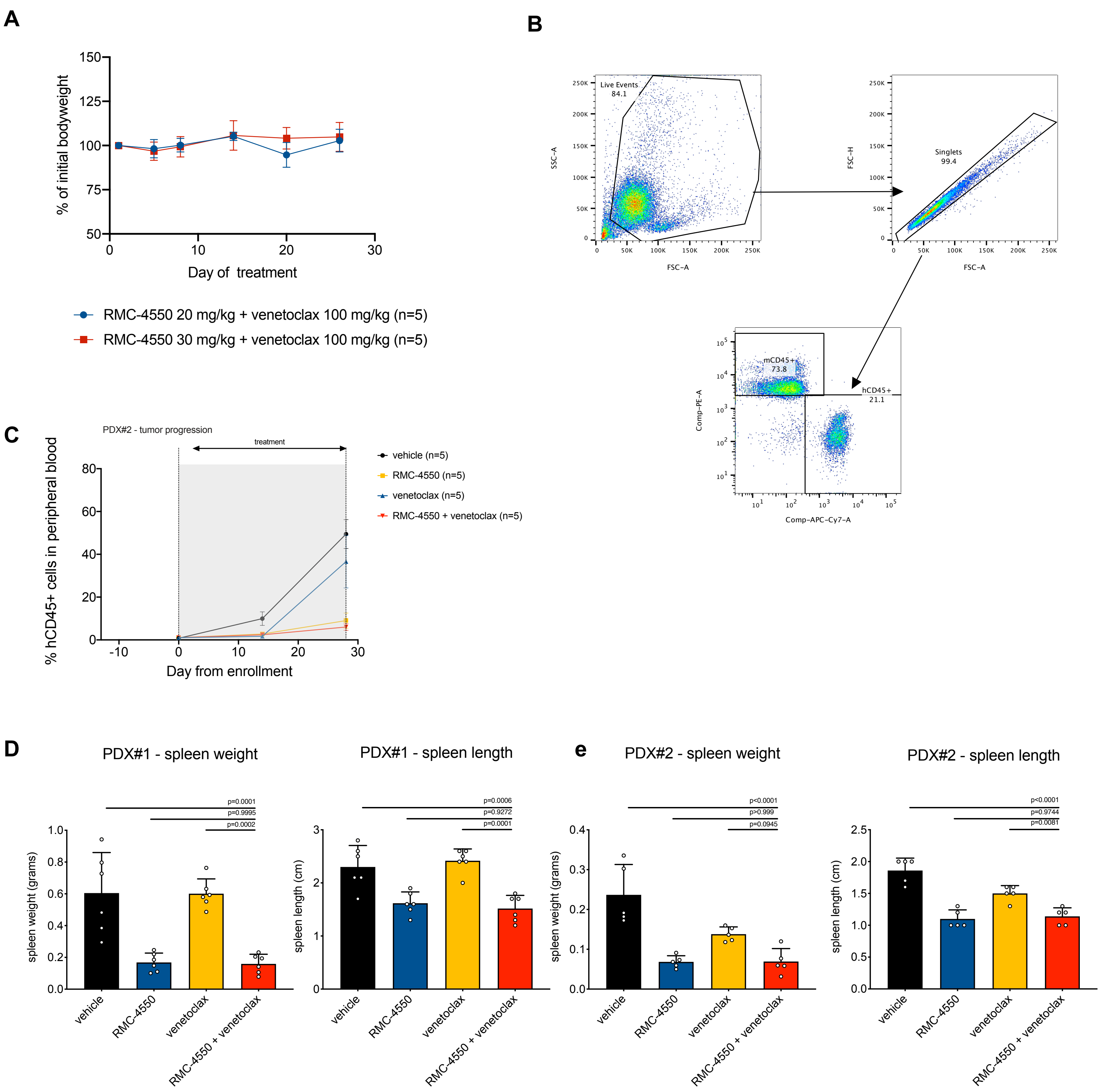

**Supplementary figure S5. Simultaneous SHP2 and BCL2 inhibition is effective in FLT3-mutant PDX AML models. A,** Body weight changes over the course of 28 days of treatment with RMC-4550 at 20 mg/kg compared to 30 mg/kg and venetoclax 100 mg/kg. Data represents mean ± SD (n=5/group). **B,** representative gating strategy for flow cytometry assessment of hCD45+ cells **C,** quantification of hCD45+ cells over the course of treatment in the four treatment groups in the PDX #2 study. **D, E,** Measurements of spleen weight and length at the study termination in PDX #1 (n=6/group, **D**) and PDX #2 – n=5/group, **E**). Data represents mean ± SD; one-way ANOVA with Tukey correction for multiple comparisons was used for statistical analysis.

#### Supplementary table S1

| Cell ilne | Molm-14 | MV4-11 | Kasumi-1 | SKNO-1 | U937 | OCIAML3 | THP-1 | NOMO-1 | HL-60 |
|---|---|---|---|---|---|---|---|---|---|
| Signaling mutation | FLT3-ITD | FLT3-ITD | KIT N822K | KIT N822K | PTPN11 G60R | NRAS Q61L | NRAS G12D | NRAS G13D | NRAS Q61L |
| RMC-4550 IC50 (nM) | 146.3 | 120 | 192.9 | 479.7 | >10000 | >10000 | >10000 | >10000 | >10000 |

#### Supplementary table S2

| *Sample* | *Local ID* | *Clinical genotype* |
|---|---|---|
| PDX #1 | HM0007 | • FLT3-ITD |
| PDX #2 | CD33CART-0005 | • FLT3-ITD<br>• NUP98-NSD1 fusion<br>• WT1 indels (in trans) |

**Supplementary table S1.** Signaling mutations and IC50 values of RMC-4550 in cell lines used for Fig. 1A data.

**Supplementary table S2.** Clinical genotypes of samples used in PDX studies.
